# Tractography density affects whole-brain structural architecture and resting-state dynamical modeling

**DOI:** 10.1101/2020.12.03.410688

**Authors:** Kyesam Jung, Simon B. Eickhoff, Oleksandr V. Popovych

**Author notes:** Corresponding author, (OP).

## Abstract

Dynamical modeling of the resting-state brain dynamics essentially relies on the empirical neuroimaging data utilized for the model derivation and validation. There is however still no standardized data processing for magnetic resonance imaging pipelines and the structural and functional connectomes involved in the models. In this study, we thus address how the parameters of diffusion-weighted data processing for structural connectivity (SC) can influence the validation results of the whole-brain mathematical models and search for the optimal parameter settings. On this way, we simulate the functional connectivity by systems of coupled oscillators, where the underlying network is constructed from the empirical SC and evaluate the performance of the models for varying parameters of data processing. For this, we introduce a set of simulation conditions including the varying number of total streamlines of the whole-brain tractography (WBT) used for extraction of SC, cortical parcellations based on functional and anatomical brain properties and distinct model fitting modalities. We observed that the graph-theoretical network properties of structural connectome can be affected by varying tractography density and strongly relate to the model performance. We explored free parameters of the considered models and found the optimal parameter configurations, where the model dynamics closely replicates the empirical data. We also found that the optimal number of the total streamlines of WBT can vary for different brain atlases. Consequently, we suggest a way how to improve the model performance based on the network properties and the optimal parameter configurations from multiple WBT conditions. Furthermore, the population of subjects can be stratified into subgroups with divergent behaviors induced by the varying number of WBT streamlines such that different recommendations can be made with respect to the data processing for individual subjects and brain parcellations.

**Author summary:** The human brain connectome at macro level provides an anatomical constitution of inter-regional connections through the white matter in the brain. Understanding the brain dynamics grounded on the structural architecture is one of the most studied and important topics actively debated in the neuroimaging research. However, the ground truth for the adequate processing and reconstruction of the human brain connectome *in vivo* is absent, which is crucial for evaluation of the results of the data-driven as well as model-based approaches to brain investigation. In this study we thus evaluate the effect of the whole-brain tractography density on the structural brain architecture by varying the number of total axonal fiber streamlines. The obtained results are validated throughout the dynamical modeling of the resting-state brain dynamics. We found that the tractography density may strongly affect the graph-theoretical network properties of the structural connectome. The obtained results also show that a dense whole-brain tractography is not always the best condition for the modeling, which depends on a selected brain parcellation used for the calculation of the structural connectivity and derivation of the model network. Our findings provide suggestions for the optimal data processing for neuroimaging research and brain modeling.

## Introduction

Some 15 years ago, the human brain connectome was introduced to understand functional brain states which are emerged by structural architecture [1]. Over more than a decade, researchers have been investigating the human connectome to elucidate the relationship between structure and function [2–5]. Recently, network neuroscience provides integrative perspectives to validate biophysically realistic models via structural connectome [6]. However, the lack of ground truth and golden standards for the calculation of the human connectome caused a central body of ongoing debates in the literature to validate the macroscopic structural and functional connectivity from neuroimaging data of the human brain [7–9]. In addition, no consensus method has been accepted so far as a standardized approach for calculating the whole-brain connectome [10,11]. Many studies have investigated the effects of the data processing on the obtained results with respect to reproducibility with different methodologies for structural architecture [12,13], functional homogeneity [14,15], and cortical resolutions for brain modeling [16]. At this stage, researchers summarized the influence of data processing for structural brain network measures [17]. Nevertheless, most of the used techniques, algorithms, and parameters for processing the neuroimaging data remain at the level of the best practice lacking a solid theoretical foundation.

Without the ground truth, a model-based approach can be a possible way to investigate the impact of the data processing on the observed brain dynamics and reveal the corresponding mechanisms [18]. At this, it is assumed that the considered mathematical models derived from the interactions between brain regions can closely simulate the dynamics of the brain responses. By comparing the simulated and empirical data, we can address the model performance as given by the results of the model fitting and thoroughly explore the model parameters and dynamics. Consequently, we can apply the model validation to evaluate the data processing by searching for the optimal model parameters that provide the best fitting of the model against the empirical data [19–21]. Such an evaluation procedure can be repeated for several modeling conditions, where the parameters of the data processing are varied. In this manner, we can systematically approach the optimal modeling condition and data parameters used for the data processing, which enhances the agreement between the simulated and empirical data.

Previous studies have used different whole-brain tractography (WBT) densities ranging from 5K to 3M tracked streamlines for the human connectome [16,22–25]. However, the impact of the WBT density on the human connectome is still unclear. Besides, the derivation of the whole-brain models essentially relies on the underlying network calculated from the whole-brain empirical structural connectivity (SC) that provides the brain architecture serving as a backbone of the brain dynamics [19–21, 23]. Due to the lack of the ground truth and standardized data processing for the whole-brain SC-networks, it is difficult to evaluate whether the selected parameters of the data processing for WBT density (e.g., the number of WBT streamlines) are reliably reflecting the brain architecture, and what are the optimal values for modeling (similarity between simulated and empirical data). In this study, we address the latter problem and search for the optimal configurations which could lead to the optimal SC extraction resulting in the best fit between the simulated and empirical data.

The broad spectrum of the computational models used for simulation of the brain dynamics ranges from the micro- to the macro-scale [21,26–30]. Besides the computational modeling concepts, the responses of brain regions can be considered as a harmonized signal [31]. Thus, we can also use simple mathematical models of coupled oscillators to describe oscillating brain activity [32–34]. In particular, systems of coupled phase and generic limit-cycle oscillators were suggested by previous studies for modeling cortical oscillations of the resting-state blood oxygen level-dependent (BOLD) dynamics [19,33,35–37]. In this study, we consider such a system of coupled phase oscillators to model the slow oscillations of the resting-state BOLD dynamics.

The main topic of the current study is to investigate the impact of the WBT streamline number used for calculation of SC and the average streamline path-length (PL) between brain regions on the simulation results. We considered a system of coupled phase oscillators with delayed coupling [38], where the anatomical information about brain structural architecture (SC and PL) from diffusion-weighted MRI (dwMRI) was used for its derivation, i.e., to build the model network and approximate the coupling weights and time delay between the network nodes. The latter are the brain regions parceled according to a given brain atlas, and we consider two distinct brain parcellations based on anatomical and functional brain properties. We systematically explored the model parameter space of two free parameters of global coupling and global delay. We also used two model fitting modalities as given by 1) similarity (Pearson’s correlation) between simulated and empirical functional connectivity (FC) as a goodness-of-fit of the model and 2) similarity between simulated FC and empirical SC to probe the dynamics of the model as related to its structural network. The obtained simulation results were compared with each other across subjects and simulation conditions, which allowed us to scrutinize the effects of structural architecture modulated by varying WBT density and brain parcellations on the model validation. The used approach can also lead to a better understanding of the properties of the obtained data influenced by selected data processing, which can play a key role for the brain modeling as well as data analytics.

## Results

The workflow of the current study schedule is illustrated in Fig 1. To evaluate the impact of the WBT density as given by the total number of WBT streamlines and the two considered brain atlases on the quality of the model validation, we first investigated how the properties of the structural connectome get affected when the WBT resolution varies, which might influence the dynamics of the whole-brain models. Then the distributions of the optimal parameters and the maximal similarities between simulated and empirical data were studied for the considered simulation conditions. We applied three criteria for the subject stratification, which provides an insight into the impact of the WBT density on the model performance for individual subjects and suggests optimal configurations of the data processing parameters.

**Fig 1.**
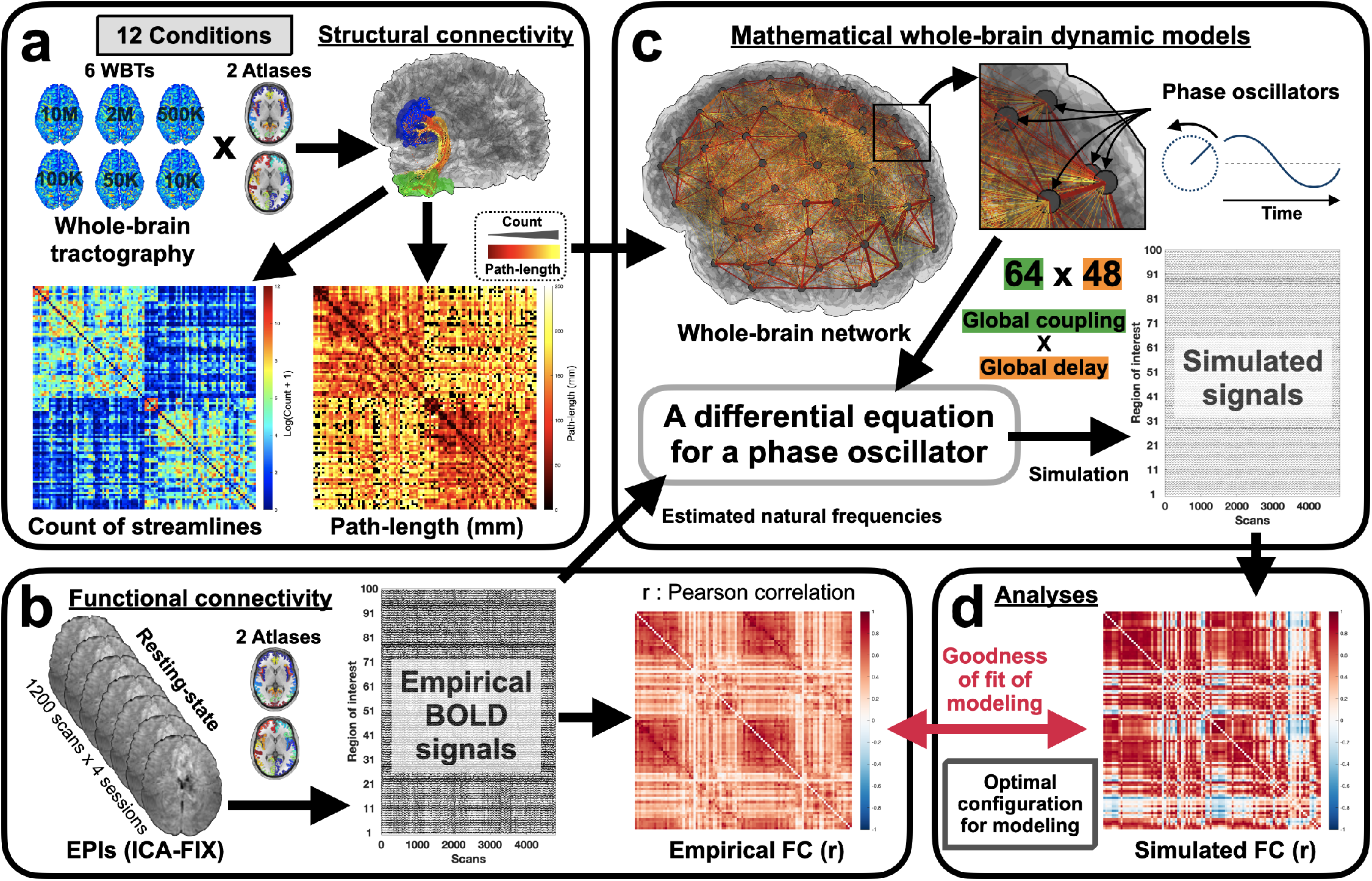
Workflow of the current study. **(a)** The whole-brain tractography (WBT) was generated by an in-house pipeline. Structural connectivity (SC) and average path-length (PL) between brain regions were reconstructed based on a given brain parcellation/brain atlas (6 WBTs and 2 atlases). **(b)** The empirical BOLD signals were extracted for each brain region from the ICA-FIX preprocessed HCP data, and the empirical functional connectivity (FC) was calculated between BOLD signals by Pearson’s correlation coefficient. **(c)** By using the empirical SC and PL matrices, the whole-brain network was reconstructed. The network nodes representing the brain regions were equipped by the phase oscillators (Eq 1) coupled with the coupling weights (Eq 2) and time delays (Eq 3) extracted from the empirical SC and PL matrices, respectively, and with the natural frequencies extracted from empirical BOLD signals. The model generated simulated BOLD signals used for the calculation of the simulated FC. **(d)** The simulated FC compared with empirical FC and SC, and the model was evaluated by optimizing its parameters for the best correspondence/fitting between the simulated and empirical data. At this, the impact of the data processing on the model validation was evaluated and described.

### Impacts of WBT resolution on structural connectome

To evaluate the architecture of the structural networks, we calculated several main graph-theoretical network properties (see **Materials and methods**) of the empirical SC and PL of individual subjects for all considered conditions of the data processing (6 WBTs for the two atlases) and tested relationships between the network properties and the model performance at the functional model fitting. Fig 2 illustrates the similarities between SC and PL (Fig 2 a and c) and behavior of the weighted node degree, clustering coefficient, betweenness centrality, local and global efficiencies, and modularity over 6 WBT conditions (10K, 50K, 100K, 500K, 2M, and 10M streamlines) for the Schaefer atlas and the Harvard-Oxford atlas (Fig 2 b and d). The similarity of the eSC matrices to the 10M case remains relatively high except for a largest drop at 10K (Fig 2 a1 and c1). On the other hand, the PL matrices very quickly deviate from the 10M case, exhibit practically no correlation already for 100K, and are weakly anti-correlated for 10K (Fig 2 a2 and c2).

**Fig 2.**
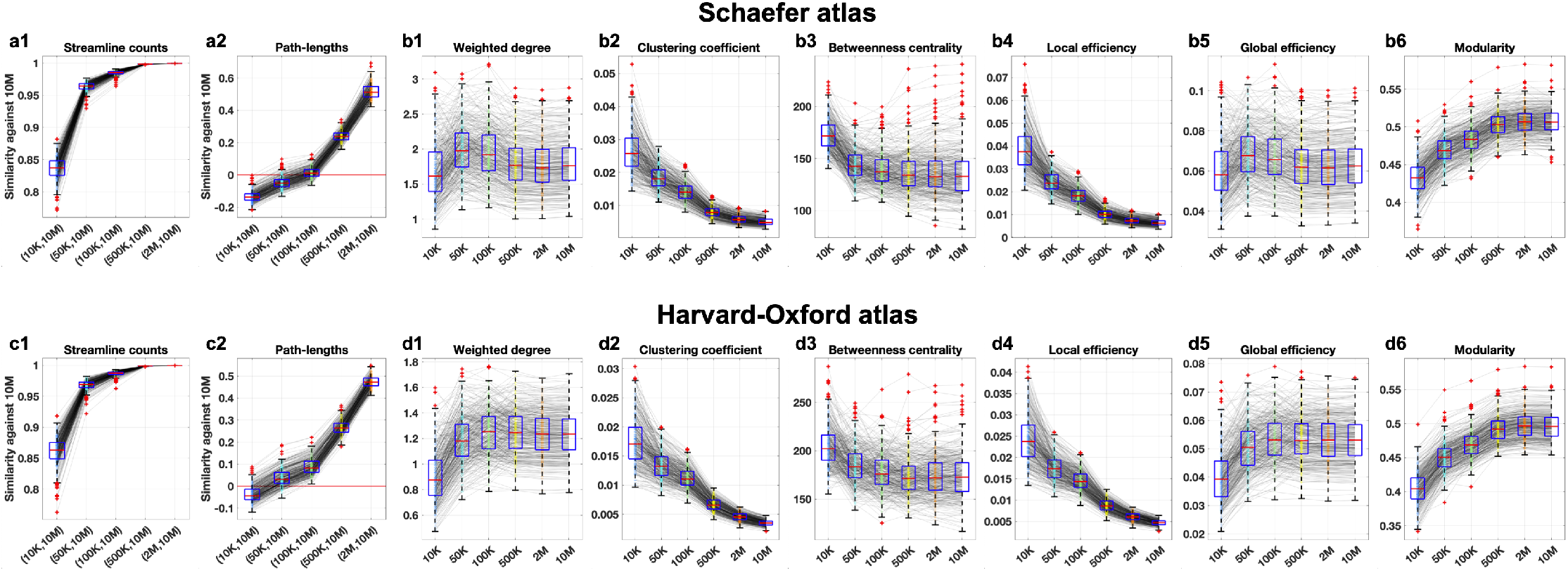
Impact of the WBT resolution on the structural architecture. Impact of the WBT resolution on the architecture of the structural networks for **(a, b)** the Schaefer atlas and **(c, d)** the Harvard-Oxford atlas. **(a, c)** Similarity of the connectivity matrices **(a1, c1)** SC and **(a2, c2)** PL against 10M case calculated by Pearson’s correlation for varying number of the WBT streamlines. **(b, d)** Variations of the network properties (1: average weighted node degree, 2: average clustering coefficient, 3: average betweenness centrality, 4: average local efficiency, 5: global efficiency, and 6: modularity) versus the streamline number. In each plot the thin gray lines depict the behavior of the illustrated quantities for individual subjects together with the box plots, where the red marks, blue boxes and red pluses indicate the medians, the interquartile ranges, and the outliers, respectively.

By increasing the number of streamlines from 10K to 10M, the number of network edges increases, and the nodes become densely connected, which results in monotonically increasing average binarized (discarded weights of edges) node degrees as expected (S1 Fig in Supplementary materials). However, the weighted node degree based on the normalized count matrices (SC divided by its mean) used in model (Eq 1) show relatively stationary behavior across the WBT conditions for dense WBT conditions (Fig 2 b1 and d1 and Table 1 WD). Similar stationary behavior can also be observed for the average betweenness centrality and the global efficiency for dense WBT conditions (Fig 2 b3, b5, d3, and d5 and Table 1 BC and GE). On the other hand, the average clustering coefficient, local efficiency and their variances decrease for dense WBT (Fig 2 b2, b4, d2, and d4 and Table 1 CC and LE), whereas the network modularity monotonically increase (Fig 2 b6 and d6). We performed a non-parametric one-way analysis of variance (Kruskal-Wallis ANOVA) test over the WBT conditions (Table 1). In summary, WBT density modulates the graph-theoretical network properties and results in similar tendencies at the group level through varying WBT resolution for both the Schaefer and the Harvard-Oxford atlases. In particular, the clustering coefficient and the local efficiency are significantly different across the WBT conditions, even between 2M and 10M (Table 1 CC and LE), where very high similarities of SC can be observed (Fig 2 a1 and c1).

**Table 1.** Sensitivity of the considered graph-theoretical network properties to the variation of the WBT density as revealed by non-parametric one-way analysis of variance (Kruskal-Wallis ANOVA) test. The corresponding p-values are presented in the right columns of the tables, where the bold p-values indicate that the respective network property significantly changes (Bonferroni corrected *p* < 0.05) when the number of WBT streamlines varies in the range indicated in the left columns of the tables. The results are shown for the Schaefer atlas (upper table) and the Harvard-Oxford atlas (lower table), and the abbreviations in the upper rows denote the network properties. WD: average weighted node degree, CC: average clustering coefficient, BC: average betweenness centrality, LE: average local efficiency, GE: global efficiency, and MQ: modularity Q.

| Schaefer atlas | WD | CC | BC | LE | GE | MQ |
| --- | --- | --- | --- | --- | --- | --- |
| 10K, 50K, 100K, 500K, 2M, 10M | <0.001 | <0.001 | <0.001 | <0.001 | <0.001 | <0.001 |
| 50K, 100K, 500K, 2M, 10M | <0.001 | <0.001 | <0.001 | <0.001 | <0.001 | <0.001 |
| 100K, 500K, 2M, 10M | <0.001 | <0.001 | 0.009 | <0.001 | <0.001 | <0.001 |
| 500K, 2M, 10M | 0.994 | <0.001 | 0.920 | <0.001 | 0.999 | 0.011 |
| 2M, 10M | 0.916 | <0.001 | 0.929 | <0.001 | 0.947 | 1.000 |
| Harvard-Oxford atlas | WD | CC | BC | LE | GE | MQ |
| 10K, 50K, 100K, 500K, 2M, 10M | <0.001 | <0.001 | <0.001 | <0.001 | <0.001 | <0.001 |
| 50K, 100K, 500K, 2M, 10M | <0.001 | <0.001 | <0.001 | <0.001 | <0.001 | <0.001 |
| 100K, 500K, 2M, 10M | 0.992 | <0.001 | 0.012 | <0.001 | 1.000 | <0.001 |
| 500K, 2M, 10M | 0.996 | <0.001 | 1.000 | <0.001 | 1.000 | 0.005 |
| 2M, 10M | 1.000 | <0.001 | 1.000 | <0.001 | 1.000 | 0.913 |

### Impacts of WBT resolution on model fitting

The system of coupled phase oscillators (Eq 1) was simulated with varying the global delay τ and the global coupling *C*, and the similarity (the two model fitting modalities) between sFC and empirical data (eFC and eSC) was calculated and assigned to that parameter value, which resulted in (*C, τ*)-parameter planes of the model fitting modalities between simulated and empirical data. Fig 3 shows the obtained parameter planes and the distributions of the optimal parameters over all subjects and simulated conditions (6 WBTs and the Schaefer and the Harvard-Oxford atlases) for the two fitting modalities (sFC versus eFC and sFC versus eSC). The maximal goodness-of-fit between sFC and eFC was observed for small delays for both atlases, see Fig 3 a-d, where the red dots depicting large similarity values are concentrated at the left side of the parameter plane demonstrating, however, different cluster shapes for the Schaefer and the Harvard-Oxford atlases. We also note here that the latter atlas could lead to a stronger fit between the sFC and eFC, compare Fig 3 a7 and c7. In contrast, in the case of the structure-functional model fitting between sFC and eSC (Fig 3 e-h), both atlases demonstrate a similar range of the correspondence (correlation) between simulated and empirical data, however, the maximal similarity can also be attained for large delay (Fig 3 f and h).

**Fig 3.**
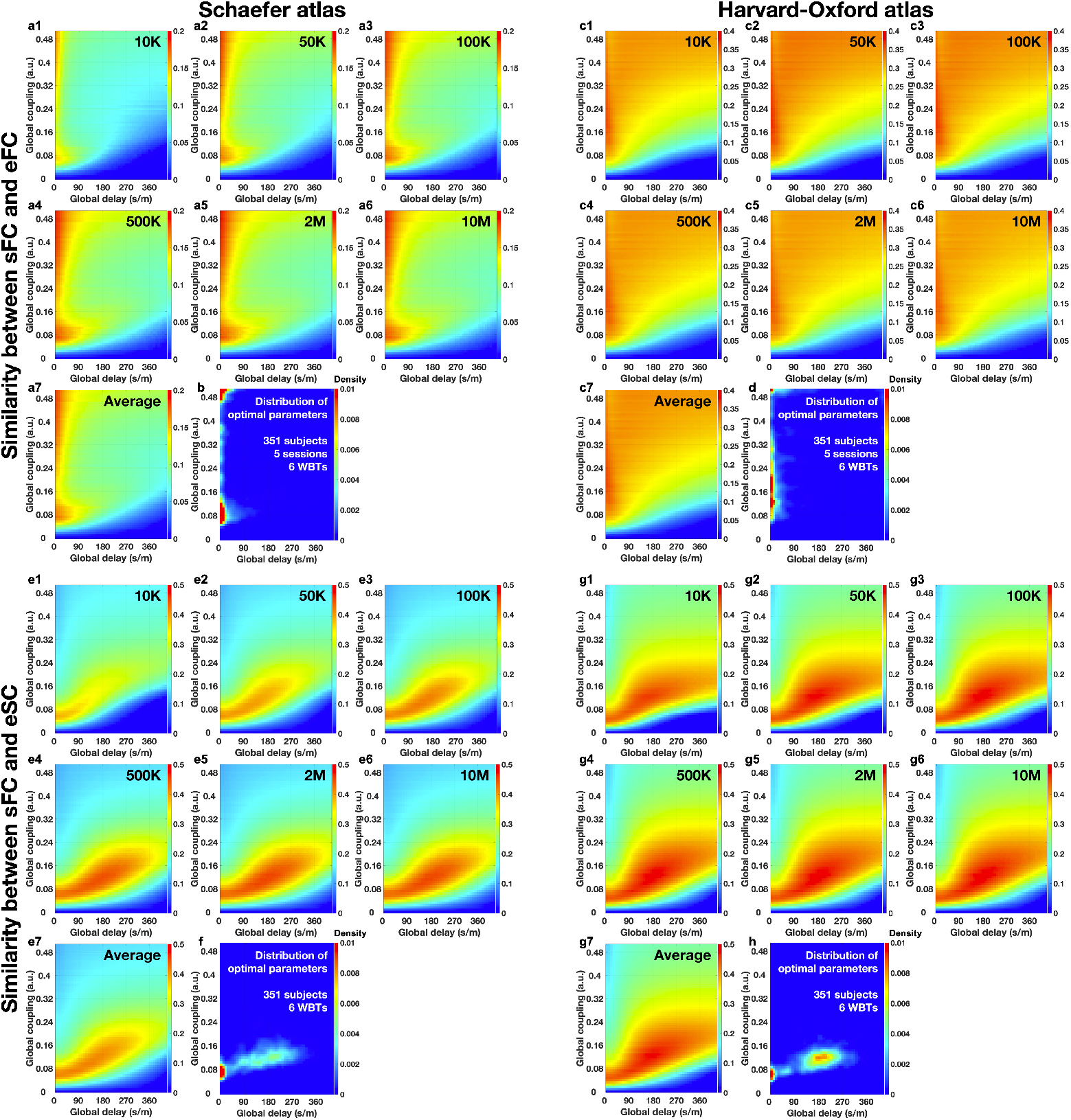
Parameter planes and the distributions of the optimal model parameters (*C, τ*) for the two model fitting modalities between simulated and empirical data. Parameter planes are averaged **(1-6)** over all subjects (n = 351) separately for any of the simulation conditions (WBT resolutions and atlases) and **(7)** also over all considered WBT resolutions as indicated in the plots. The correspondence between the simulated and empirical data was calculated between **(a-d)** simulated FC and empirical FC and **(e-h)** simulated FC and empirical SC for **(a, b, e, f)** the Schaefer atlas and **(c, d, g, h)** the Harvard-Oxford atlas. The Pearson correlation between the connectivity matrices is depicted by color ranging from small (blue) to large (red) values. **(b, d, f, h)** Distributions of the optimal model parameters of the best model fitting calculated for all individual subjects and simulation conditions.

During the model validation for individual subjects under the 12 considered conditions (6 WBTs ×2 atlases), we also searched for the optimal model parameter, where the maximal similarity between sFC and empirical data (eFC and eSC) was achieved. The distributions of the optimal parameters are depicted in Fig 3 b, d, f, and h for the two fitting modalities and the two brain atlases. In agreement with this results, the best fit between sFC and eFC is attained for small delays (Fig 3 b and d), whereas the strongest structure-function correspondence between sFC and eSC can also be observed for large delays (Fig 3 f and h). In the latter case, the parameter distributions apparently demonstrate a two-cluster shape of small and large delays, which is addressed in detail below.

Together with the optimal model parameters for individual subjects, we also collected the corresponding maximal similarities between the simulated and empirical data, which are illustrated in Fig 4 for the 12 simulated conditions and for the two fitting modalities of the correspondence between sFC and eFC (Fig 4a) and between sFC and eSC (Fig 4b). Results of the functional model fitting in all conditions (Fig 4a) were not from the normal distributions, where the null hypothesis was rejected by χ^2^ goodness of fit test with p < 0.05. In the case of the structure-functional model fitting (Fig 4b), the conditions of 500K, 2M, and 10M WBT streamlines for the Schaefer atlas and 100K, 500K, 2M, and 10M streamlines for the Harvard-Oxford atlas were also not from the normal distributions. Therefore, Kruskal-Wallis test was used for testing significant difference in all conditions. Consequently, we performed Wilcoxon signed rank one-tail test to evaluate whether the maximal similarities between the simulated and empirical data for one condition are significantly higher or lower than those for the other conditions (see p values in Fig 4). For the functional model fitting (sFC versus eFC) and the Schaefer atlas (Fig 4a, blue violins), the models with 2M and 10M WBTs performed better than with the other WBTs, and the performance of the model decreased when the number of streamlines decreased. On the other hand, the functional model fitting for the Harvard-Oxford atlas revealed the optimal condition at 50K or 100K WBT (Fig 4a, orange violins). Furthermore, the model could fit better to eFC for the Harvard-Oxford atlas, which was also observed in Fig 3. For the structure-functional model fitting (sFC versus eSC), the situation is different, where 2M or 10M WBTs are preferable for the strongest correspondence between the simulated and empirical data for both atlases demonstrating approximately similar extent of the maximal model fitting (Fig 4b, see also Fig 3).

**Fig 4.**
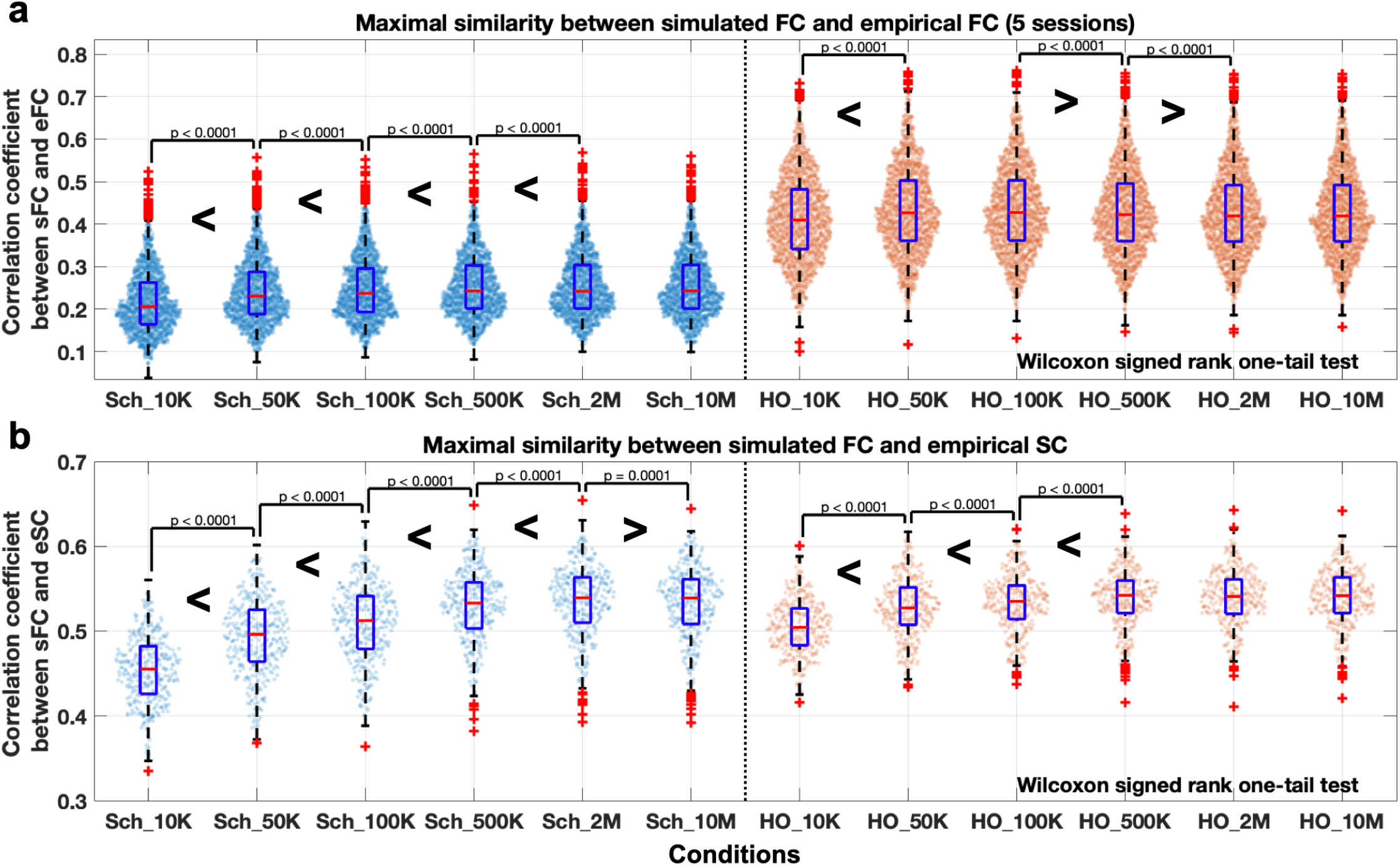
Results of the model fitting to the empirical data versus 12 simulation conditions (6 WBTs × 2 atlases). The distributions of the maximal similarities for individual subjects between **(a)** simulated FC and empirical FC and **(b)** simulated FC and empirical SC are shown as violin plots for 12 conditions indicated on the horizontal axes for the Schaefer atlas (blue violins) and the Harvard-Oxford atlas (orange violins). The results of the pairwise comparisons between the conditions (Wilcoxon signed rank one-tail test) are also indicated with the corresponding p-values in the cases of statistically significant differences (Bonferroni corrected p < 0.05). Notation “SCH” on the horizontal axes is for the Schaefer atlas, and “HO” is for the Harvard-Oxford atlas, for instance, “SCH_10M” stands for the condition of Schaefer atlas with 10M streamlines. For the box plots the red marks, blue boxes and red pluses indicate the medians, the interquartile ranges, and the outliers, respectively.

### Relationships between network properties and the functional model fitting

As discussed above, the WBT density modulates the structural connectome. Consequently, it can also influence the dynamics of the model (see parameter planes over the simulation conditions in Fig 3). In this section, we investigate the effects of the graph-theoretical network properties modulated by WBT density on the model performance.

For each of the considered 10 network properties, we tested the relationships between their values and the maximal similarity between sFC and eFC as given by the Pearson’s correlation across 6 WBT conditions for each individual subject. The considered network properties demonstrate a pronounced agreement with the maximal goodness-of-fit values at the level of individual subjects (Fig 5 a1 and b1), where the distributions of the correlation coefficients are significantly shifted from zero (except for the averages of weighted node degree, the standard deviations of betweenness centrality, and global efficiencies for the Schaefer atlas and the standard deviations of betweenness centrality for the Harvard-Oxford atlas). Furthermore, this relationship is reproducible to retest over individual 5 sessions (see S2 Fig in Supplementary material). Based on the results illustrated in Fig 5 a1 and b1 and S2 Fig, we can conclude that the changes in the model performance for the individual subjects are related to the changes in the network properties across different WBTs.

**Fig 5.**
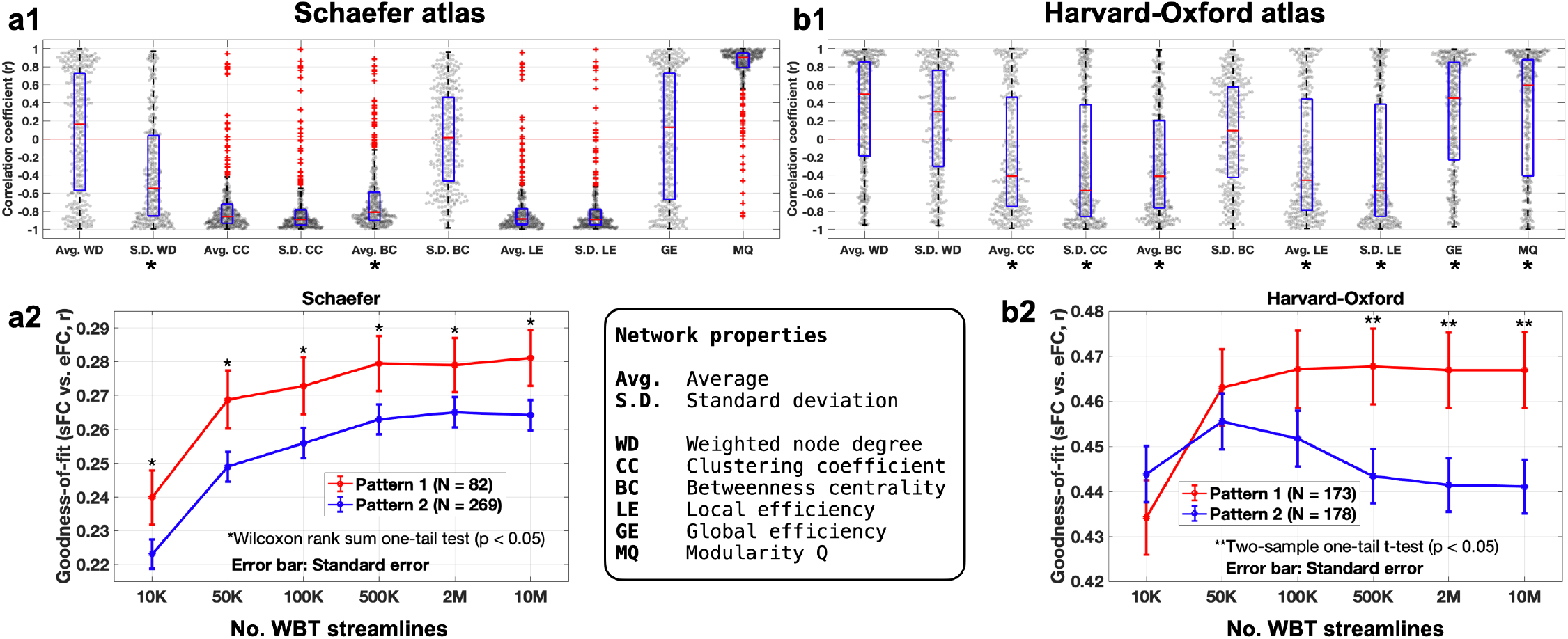
Relationships between the network properties and the results of the functional model fitting. Relationships between the network properties and the results of the functional model fitting (maximal similarity between sFC and eFC) for individual subjects for **(a)** the Schaefer atlas and **(b)** the Harvard-Oxford atlas. **(a1, b1)** Distributions of the Pearson’s correlation coefficients calculated across 6 WBT conditions for individual subjects between a given network property indicated on the horizontal axes and the maximal goodness-of-fit values. The gray dots represent the values for individual subjects, and the box plots illustrate the medians (red lines), the interquartile ranges (blue boxes) and the outliers (red pluses). The asterisks below the x-axes labels indicate statistically significant differences in the maximal goodness-of-fit values between the two subgroups of subjects with positive and negative correlations (*p* < 0.05 of two-sample one-tail t-test). **(a2, b2)** The results of the functional model fitting versus different numbers of the WBT streamlines for the two subject subgroups of pattern 1 and pattern 2 as indicated in the legends based on the statistically significant split of the subjects for the network properties marked by asterisks in plots a1 and b1, see the **Materials and methods** and text for details. The error bars indicate the standard error, and the asterisks denote the simulation conditions, where the pattern 1 and 2 exhibit significantly different extend of the similarity between simulated and empirical data.

The distributions of the correlation coefficients for the Harvard-Oxford atlas (Fig 5 b1) less deviate from zero than those for the Schaefer atlas (Fig 5 a1) indicating a complex relationship between the network properties and the maximal goodness-of-fit of the model. To address this relationship in more detail, for every considered network metrics, we split the subjects into two subgroups of positive or negative correlation across the WBT conditions between a given network property and the maximal goodness-of-fit values, see also S3 Fig and S4 Fig in Supplementary materials for more details. Subject subgroups with the higher goodness-of-fit values were intersected for the network metrics marked by asterisks in Fig 5 a1 and b1 with significant difference between the subgroups. The obtained intersection formed the stratified subject group referred to as pattern 1, whereas the remaining subjects are united into pattern 2 as mentioned in the **Materials and methods**, see also S5 Fig in Supplementary materials.

Based on the results of the tests, for the Schaefer atlas, we selected subjects exhibiting positive correlation with the standard deviation of weighted node degree (S.D. WD+) and negative correlation with the average betweenness centrality (Avg. BC-) for pattern 1, which have significantly higher values of goodness-of-fit of the model than those of the complementing subgroups (S.D. WD- and Avg. BC+), respectively. The intersection of the two selected subgroups, i.e., S.D. WD+ (n = 93) ⋂ Avg. BC- (n = 329) = 82, constituted the stratified pattern 1, whereas the rest of the subjects (n = 269) were grouped into pattern 2. For the Harvard-Oxford atlas, we selected subjects from the intersection of the following subgroups derived as above of positive and negative correlations with the network metrics, which showed significantly higher goodness-of-fit values than the complementing subgroups: Avg. CC-, S.D. CC-, Avg. BC-, Avg. LE-, S.D. LE-, GE+, and MQ+ (see the rounded box in Fig 5 for abbreviations). As above, the sign “+” or “-” after the property name indicates the corresponding subgroups of subjects exhibiting positive or negative correlations with the considered network properties, respectively. Such an intersection of the subgroups resulted in a stratified pattern 1 containing 173 subjects complemented by the others, i.e., 178 subjects into pattern 2. We applied the pattern 1 and 2 splitting for each atlas for the first criterion of the stratification analysis.

We found that the two patterns of the split subjects subgroups demonstrate significantly different quality of the goodness-of-fit of the model depending on the WBT conditions (Fig 5 a2 and b2). We also found that the patterns 1 and 2 exhibit different behavior of the maximal goodness-of-fit values when the WBT resolution varies. In particular, for the Schaefer atlas the quality of the model validation monotonically increases for both patterns for higher WBT resolution (Fig 5 a2), whereas for the Harvard-Oxford atlas, the pattern 2 apparently demonstrates a non-monotonic behavior with an optimal point at 50K of the WBT streamlines (Fig 5 b2). χ^2^ goodness of fit test was used to test for a normal distribution for each condition of pattern 1 and pattern 2, and the Wilcoxon rank sum one-tail test was then used for a non-parametric test of the difference between the patterns if it is rejected by the χ^2^ test. Otherwise, two-sample one-tail t-test was used for comparing normal distributions of pattern 1 and pattern 2. We also tested the changes of the goodness-of-fit of the model for each pattern when the WBT resolution varies by using Wilcoxon signed rank test. As a result, for the Schaefer atlas, 500K or more streamlines of the pattern 1 and 2M or more streamlines of the pattern 2 showed significantly higher goodness-of-fit values than for the sparser WBT conditions. For the Harvard-Oxford atlas, 100K or more streamlines of the pattern 1 showed significantly higher goodness-of-fit values than sparser WBT conditions. However, 50K streamlines of the pattern 2 is the optimal condition that shows significantly higher correspondence between the simulated and empirical data than for the other conditions.

Based on the presented results, we can conclude that the optimal number of the WBT streamlines should be considered large (~500K-10M) for the Schaefer atlas (Fig 5 a2). Interestingly, the best goodness-of-fit of the model for the Harvard-Oxford atlas can be reached for much sparser WBT at ~50K streamlines for more than 50% of the subjects (Fig 5 b2).

### Effects of time delay on model validation

Based on the clustered distributions of the optimal model parameters of the maximal structure-function similarity between sFC and eSC (Fig 3 f and h), we divided the optimal parameter points and the corresponding subjects into two clusters (Fig 6). In such a way, the cluster of parameter points with small delay (cluster 1) was split from the other points characterized by relatively large delay (cluster 2) based on their bimodal distributions (Fig 6, the red dotted lines in the histograms in the bottom plots). By dividing the subjects into the two subgroups corresponding to the above clustering of their optimal parameters, we found that the maximal goodness-of-fit values of the functional model fitting are significantly higher in cluster 2 than in cluster 1 consistently for all simulation conditions (all WBTs and both atlases), see Fig 6 (upper plots). Similar effects can also be observed for the structure-functional model fitting between sFC and eSC (see S6 Fig a2 and b2 in Supplementary materials), where the time delay in the coupling played a constructive role in the model validation against empirical data.

**Fig 6.**
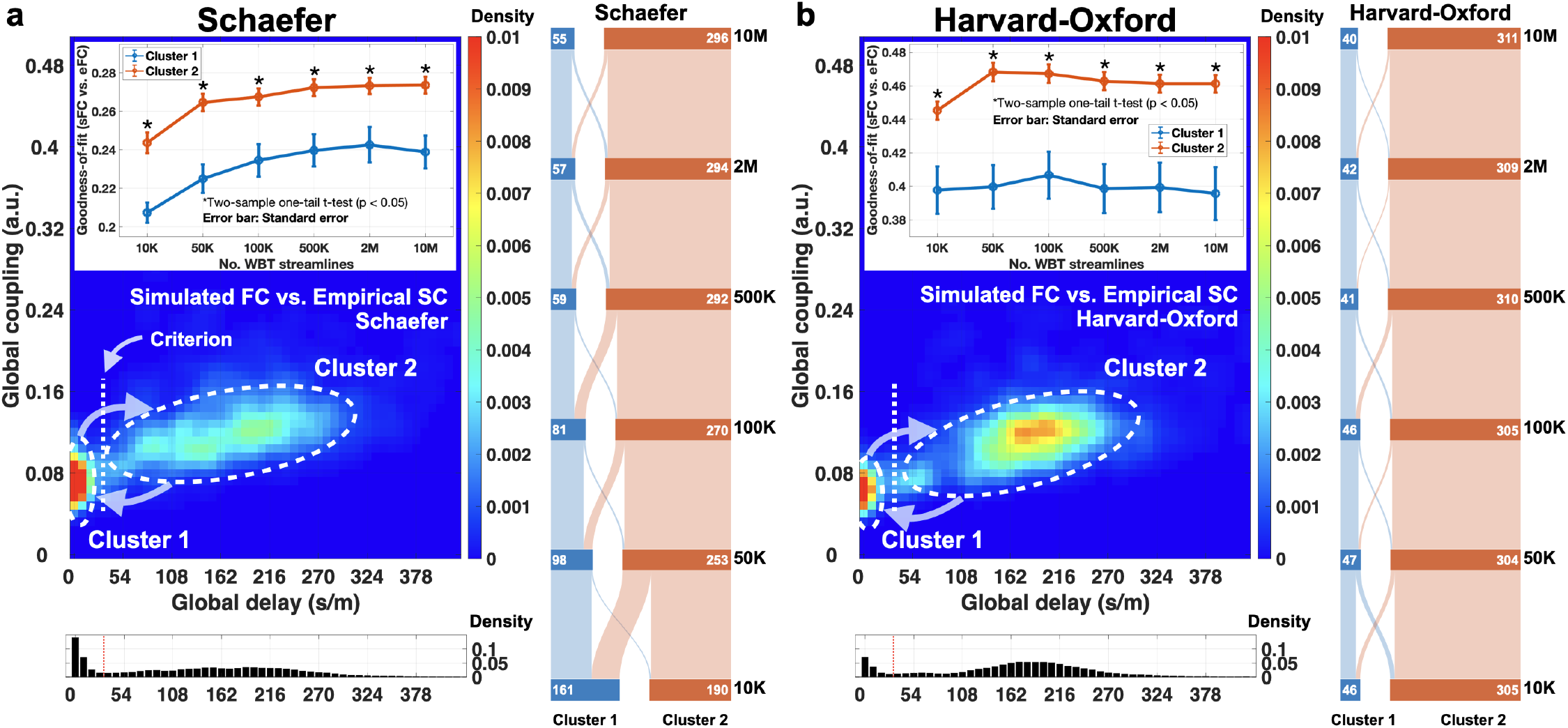
Clusters of the optimal model parameters of the maximal similarity between simulated FC and empirical SC. The optimal parameters for **(a)** the Schaefer atlas and **(b)** the Harvard-Oxford atlas from Fig 3 f and h, respectively, (n = 2106 values for 351 subjects and 6 WBTs) were split into two subgroups as illustrated in the two lower plots, where the one- and two-dimensional distributions of the optimal parameters are depicted. The upper plots with error bars show the maximal similarity of the functional model fitting between simulated FC and empirical FC of the concatenated fMRI session for the subjects from the two clusters versus the number of the WBT streamlines. The alluvial plots to the right schematically illustrate the interchange of the cluster members when the number of streamlines varies from 10M to 10K. The white numbers in each WBT step indicate the number of subjects in the clusters.

These results also establish a connection between the two fitting modalities and the time delay, where the impact of the latter was not observed in the distributions of the optimal parameters of the functional similarity between sFC and eFC (Fig 3 b and d) and can only be revealed by mediation of the structure-function correspondence. The broad distributions of global delays in cluster 2 can also be related with the natural frequencies of the phase oscillators (Eq 1). As mentioned in the **Materials and methods**, we used natural (intrinsic) frequencies for the individual oscillators estimated from the empirical BOLD signals in the range from 0.01 Hz to 0.1 Hz. The mean natural frequency of the model is also varying across subjects, and we found a well-pronounced negative correlation between the natural frequencies and the optimal delays for the similarity between sFC and eSC (see S7 Fig and S8 Fig in Supplementary materials).

When the number of the WBT streamlines varies, subjects may exchange their membership in the two clusters (Fig 6, the vertical alluvial plots). Interestingly, for the Schaefer atlas, the ratio of subjects in the two clusters is gradually changing when WBT is getting sparser (from 10M to 10K), where more and more subjects move to cluster 1 approximately balancing the subgroup sizes at 10K case (Fig 6a, the alluvial plot). In contrast, there are only small exchanges of the subject members between clusters for the Harvard-Oxford atlas keeping the group sizes approximately constant for all WBT conditions (Fig 6b, the alluvial plot), where cluster 2 contains most of the subjects as is for the Schaefer atlas for the case of 10M of the WBT streamlines. We used the splitting of the subjects into the discussed two clusters as the second criterion of the stratification analysis.

### WBT-induced changes of model performance

In the previous sections we observed that the behavior of the maximal goodness-of-fit versus the WBT conditions is not akin to each other for different atlases. For example, for the Schaefer atlas, the maximal similarity between sFC and eFC monotonically increases when the WBT is getting denser (Fig 5 a2), and the corresponding curve had a positive slope. On the other hand, the maximal goodness-of-fit for the Harvard-Oxford atlas may exhibit a non-monotonic behavior and attained the maximal values at 50K WBTs (Fig 5 b2). We therefore explicitly searched for such a divergent dynamics, looked for the subjects with the best model performance for the most sparse or the most dense WBT, and divided the subjects into two subgroups based on the behavior of the model performance when the number of WBT streamlines varies. In this way, we compared the maximal goodness-of-fit values of the functional model fitting for the two extreme cases of 10K and 10M streamlines and separate the subjects if the behavior of the model fitting results demonstrated opposite tendencies. We thus distinguished positive and negative slopes of the maximal goodness-of-fit of the model versus the WBT conditions, when the latter increases from 10K to 10M. Fig 7 illustrates the different dynamics of the maximal goodness-of-fit of the two subgroups of the subjects for the two atlases. For the Harvard-Oxford atlas, the both subgroups contain large fractions of the entire subject population with the positive slope (n = 248) and the negative slope (n = 103) (Fig 7b). On the other hand, the subjects split very unevenly for the Schaefer atlas, and most of them (n = 339) exhibited positive slope, where the similarity between simulated and empirical data monotonically increases when the number of streamlines increases (Fig 7a).

**Fig 7.**
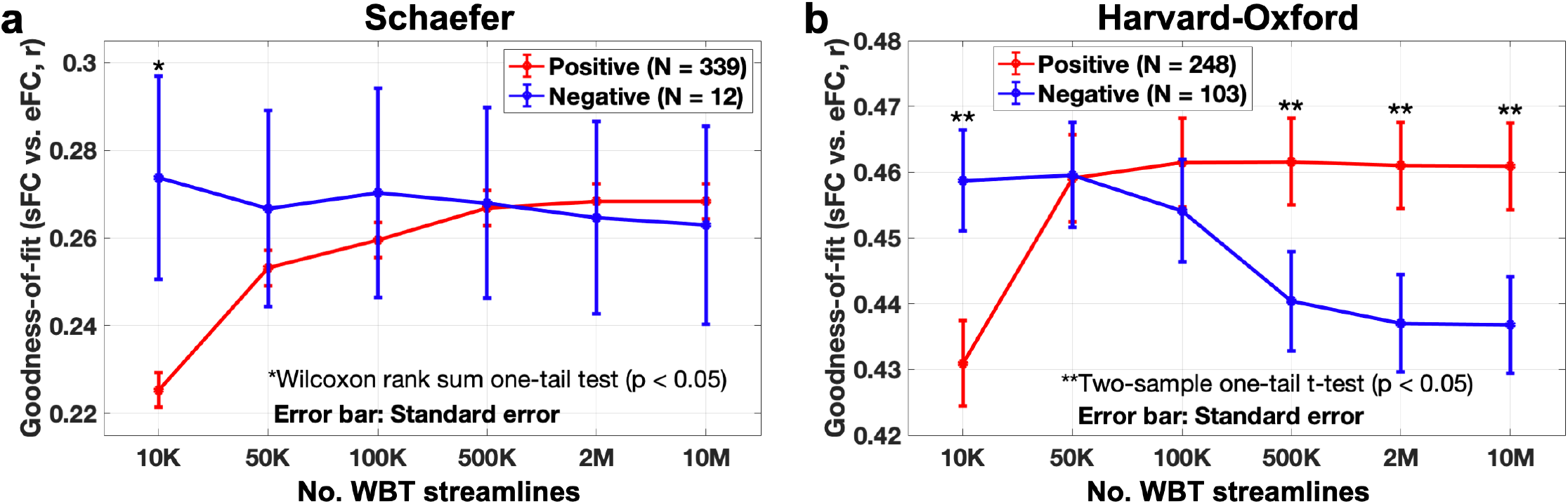
Subject stratification according to the model performance across 6 WBTs. **(a, b)** The maximal goodness-of-fit of the functional correspondence between simulated FC and empirical FC for **(a)** the Schaefer atlas and **(b)** the Harvard-Oxford atlas and for the two groups of the subjects stratified according to the third criterion (see **Variation of the model performance** in the **Materials and methods**). The latter is based on the behavior (positive/negative slopes) of the maximal similarity versus the WBT conditions (see text for details) as indicated in the legends, where the number of subjects in each group is also pointed out. The asterisks indicate the statistically significant differences between the two subject groups (p < 0.05, two-sample one-tail t-test for normal distributions and Wilcoxon rank sum one-tail test for non-parametric test).

Each split subgroup was tested for a normal distribution by χ^2^ goodness of fit test over WBT densities. The null hypothesis of the χ^2^ test for the Schaefer atlas was rejected for each subgroup and each condition, however, in the case of the Harvard-Oxford atlas, the null hypothesis was not rejected. Consequently, we performed Wilcoxon signed rank test for the Schaefer atlas and two-sample paired t-test for the Harvard-Oxford atlas. As a result, for the Schaefer atlas, the subgroup with the positive slope of the 2M or more streamlines WBT conditions showed significantly higher goodness-of-fit of the model than sparser WBT conditions (Fig 7a, red curve). In the case of the Harvard-Oxford atlas, the subgroup with the positive slope showed significantly higher goodness-of-fit of the model with 100K or more WBT streamlines than sparser WBT conditions (Fig 7b, red curve). On the other hand, the the subgroup with the negative slope showed significantly higher goodness-of-fit of the model with 50K or less streamlines WBT conditions than denser WBT conditions (Fig 7b, blue curve). We applied the discussed approach based on the model performance as the third criterion of the stratification analysis.

### Stratification analysis

As investigated in the previous sections, the entire subject population can be split into two groups based either on the two patterns of the relationships between network properties and the functional model performance (Fig 5), the two clusters based on the distributions of the optimal parameters of the maximal similarity between sFC and eSC (Fig 6), and the behavior (positive and negative slopes) of the maximal goodness-of-fit for the correspondence between sFC and eFC versus the WBT conditions (Fig 7). By combining all three approaches, we illustrated a stratification result in the alluvial plots in Fig 8, which show the proportions of the stratified subjects when the above stratifying criteria are consequently applied to the entire subject population for each atlas.

**Fig 8.**
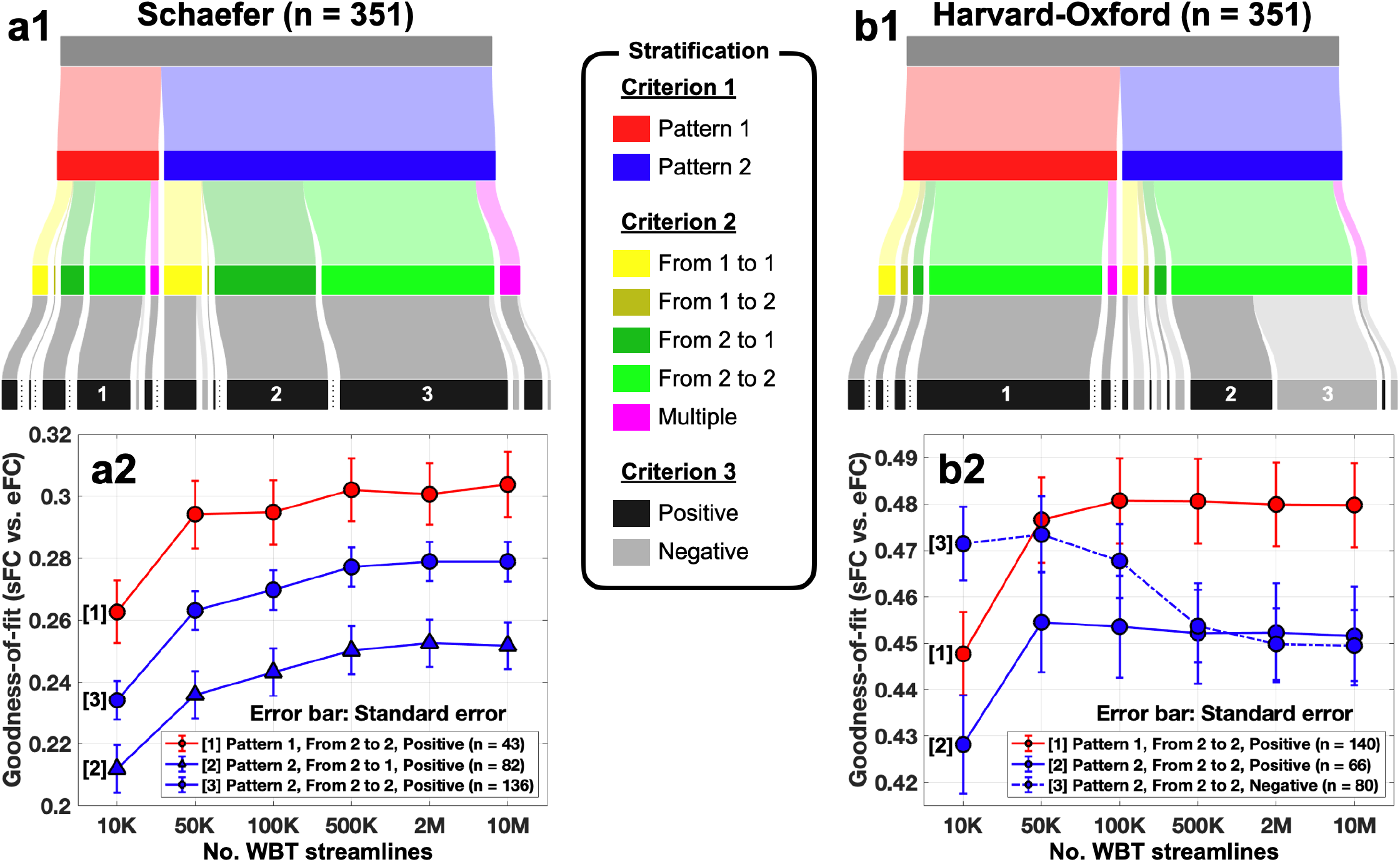
Stratification analysis with three criteria for two atlases. **(a1)** The alluvial plot shows all stratified subjects via three criteria and **(a2)** the bottom plot shows goodness-of-fits through 6 WBT conditions for large stratified groups (>35) in the case of the Schaefer atlas. **(b)** The plots by the same analyses from the Harvard-Oxford atlas.

The stratified subjects in Fig 8 a1 and b1 show different extent and behavior of the maximal goodness-of-fit values of the functional model fitting over the WBT conditions (Fig 8 a2 and b2). The first criterion provides a way to explain the changes of the fitting results across WBT densities for individual subjects involving certain properties of the empirical structural networks (Fig 5). According to the first criterion, we can expect that the subjects from pattern 2 will have the maximal model performance for sparse WBTs for the Harvard-Oxford atlas (Fig 5 b2).

The second stratification step in Fig 8 a1 and b1, reflects the interchanging behavior between the parameter clusters observed in Fig 6, which is notable for the Schaefer atlas, whereas it is not prominent for the Harvard-Oxford atlas. This is also observed in Fig 8 a1 and b1, where the overwhelming majority of subjects for the latter atlas were sorted to the group of persistent members of cluster 2, i.e., the split subgroup with large delay for the structure-functional model fitting. Finally, the third criterion practically does not differentiate the subjects into positive and negative slopes for the Schaefer atlas, which is also true for pattern 1 in case of the Harvard-Oxford atlas. However, the subjects in pattern 2 for the Harvard-Oxford atlas can still be split into two subgroups with the inclining and the declining curves of the maximal goodness-of-fit by the third criterion, which can further refine the differentiation of subjects of the best model performance at sparse WBT density (Fig 8b2, stratified groups 2 and 3, see also Fig 5).

The declining curves of the goodness-of-fit when the number of the WBT streamlines decreases imply that the optimal number of the total streamlines for the simulation with the Schaefer atlas should be considered large, for example, more than 500K: 2M or 10M of the WBT streamlines (Fig 8a2). The model evaluation with the Harvard-Oxford atlas show different optimal conditions than that for the Schaefer atlas (Fig 8b2). The optimal streamline number may depend on the stratification subgroups to which the subject belongs, and which exhibited very different behavior of the goodness-of-fit when the number of streamlines varied (Fig 8b2). For example, the optimal number of streamlines for a better model performance could range from 10M to 100K for the subjects from subgroup 1 in Fig 8b2 (solid red curve). On the other hand, for more than 20% of subjects (n = 80) of the entire subject population, i.e., for those from the stratified group 3 (Fig 8b2, dashed blue curve), the optimal conditions are at ~50K WBT streamlines, and more streamlines may lead to the degradation of the quality of the model validation. For other 18% of subjects (n = 66, group 3 in Fig 8b2, solid blue curve) a sparse WBT can also be a reasonable option.

## Discussion

The purpose of the current study was to explore how the processing of the neuroimaging data can influence the dynamics and validation of the whole-brain mathematical dynamical models informed by the empirical data. We considered several simulation conditions based on varying parameters of data processing, such as the number of total streamlines (or fibers in the brain white-matter) of WBT and brain atlases. While the latter defined how the brain is parceled into several brain regions that are considered as network nodes in the model, the former influenced the underlying SC and PL used for the calculation of the coupling weights and time delays in the coupling between nodes. We discussed how the WBT resolution can influence the structural information fed to the model and the corresponding modeling results for the considered brain atlases. We found that the parcellation with different atlases showed similar changes of the architecture of the structural networks, but distinct trends of the goodness-of-fit of the model to the empirical data across the numbers of WBT streamlines. Consequently, we suggested optimal configurations of the considered data and model parameters for the best model fit at the group level as well as for personalized models of individual subjects based on the properties of the empirical and simulated data.

The applied model-based approach followed the line of research suggested and developed in many modeling studies, see, for example, the papers [18,19,23,33,35–37] and references therein, where the potential of the whole-brain dynamical models to explain the properties of the brain dynamics and structure-function relationship was demonstrated by a detailed investigation of the correspondence between empirical and simulated brain connectomes. At this, the patterns of the underlying structural network connectivity as related to the inter-node coupling strengths and delays can play a crucial role for observing a pronounced structure-function agreement [39, 40]. It is thus important to extract the empirical SC and PL used for evaluation of parameters of the model connectivity as plausible as possible in order to obtain biologically realistic modeling results [41].

### Evaluating structural architecture for modeling

As discussed in Fig 2 and Table 1, the variation of the WBT streamline number affects the properties of the model networks calculated from the structural connectome, especially, the PL matrices, where the edges with relatively small numbers of streamlines are sensitive to reducing the total number of tracking trials. Therefore, SC extracted from relatively sparse WBT with small number of streamlines may not guarantee a higher reproducibility with stable network properties, where some edges will be disconnected or reconnected from time-to-time, when streamlines will be generated.

Within the framework of the modeling approach, the model parameters can be varied in a broad range and sense to evaluate their impact on the simulated dynamics. As related to the discussed network topology, beyond the variation of the global coupling strength, the network edges approximating the anatomical connections between brain regions can be removed to obtain a better fit between simulated and empirical FC of schizophrenia patients [42]. Aiming at the best correspondence between simulated and empirical data, new inter-region anatomical connections were allowed to be created, or existing structural connections to be rewired according to algorithms based on the differences between the simulated and empirical FC including the gradient-descent method [43, 44]. The model connectivity can be composed of both empirical SC extracted from dwMRI data and local intra-cortical connections incorporated into the model based on the distance-dependent approximations [16]. Among many possible ways of SC variation for the best model fitting, which might also require additional justifications, we propose to stay within the framework of realistically extracted signals from dwMRI data and consider the well-established approaches for the data processing. In this study, we used state-of-the-art techniques for calculation of WBT and SC [45] and investigated the impact of a constructive parameter for the structural connectome, the number of extracted streamlines on graph-theoretical measures of SC, and their influence on the modeling results.

### Effects of different parcellations on the modeling

For the extraction of the brain structural and functional connectomes and for setting up the model network, we used two paradigmatically distinct brain atlases as represented by the Schaefer atlas [46] that is based on functional MRI data, and the Harvard-Oxford atlas of anatomy-related parcellation [47] that is based on the landscape of gyri and sulci on the cortical surface. We found that the graph-theoretical properties of the structural networks built based on these two parcellations and used in the model are changing with similar tendencies across the considered WBT conditions for the both atlases (Fig 2 and Table 1). Some of the considered network properties exhibit high sensitivity to the variations of the WBT density, for example, the clustering coefficient (CC) or the local efficiency (LE), see Table 1. On the other hand, the weighted node degree (WD) or the global efficiency (GE) manifested significant changes only when the number of the calculated WBT streamlines was decreased from 10M to 100K or 50K, i.e., 100-200 times, and the sensitivity was stronger in the case of the Schaefer atlas. These findings might be of importance when the discussed network properties influence the modeling results.

We also found that the mentioned network metrics (CC and LE) with sensitive dependence on the WBT density strongly anti-correlate with the goodness-of-fit of the model for the Schaefer atlas, while the relationship is in average less pronounced for the Harvard-Oxford atlas (Fig 5 a1 and b1). Furthermore, the situation is reversed for the insensitive network properties (WD and GE), where the correlation is more enhanced for the Harvard-Oxford atlas. Given the impact of the WBT density on the properties of the structural networks (Fig 2), this may explain the clear monotonic behavior of the goodness-of-fit for the Schaefer atlas versus the number of streamlines (Fig 5a2) and apparently mixed behavior for the Harvard-Oxford atlas (Fig 5b2). In summary, some of the network metrics are characterized by different relationships with the results of the model validation for the varying number of the WBT streamlines between the parcellations, see also S3 Fig and S4 Fig in Supplementary materials for the relationships of all considered network properties. Therefore, the tractography density modulates the graph-theoretical network properties in similar changes for the considered atlases, however, it influences the dynamics of mathematical models in different ways, in particular, depending on the used brain parcellation.

### Role of time delay in the modeling

We also reported on the differences with the distributions of the optimal model parameters for the structure-functional model fitting sFC-eSC and their behavior for the two considered brain atlases, where a parameter clustering with respect to small and large delay in the coupling was observed (Fig 6). More detailed investigation revealed an apparent migration of the optimal parameter points towards the parameter cluster of small delay when the number of streamlines decreased for the Schaefer atlas. On the other hand, no such parameter (and subject) flow was detected for the Harvard-Oxford atlas, and the parameter distribution remained relatively stable for any of the considered numbers of streamlines. Such a behavior of the optimal parameters for the two considered atlases might be related to the performance of the model validation at the group level when the WBT resolution varies, which can suggest a possible mechanism associated with the impact of time delay in coupling on the model fitting results. Indeed, we observed that subjects from the parameter cluster with large delay demonstrated better quality of the model validation for both functional and structure-functional model fittings (Fig 6 and S6 Fig). In other words, if the optimal parameters for the maximal sFC-eSC correspondence have a large delay, we might expect a better correspondence between sFC and eFC. Accordingly, we might also expect that the group-averaged maximal goodness-of-fit for the Schaefer atlas will decay faster than that for the Harvard-Oxford atlas when the streamline number decreases as observed in Fig 4, because fewer optimal parameter points with large delay can be found for a sparser WBT for the former atlas. It is also important to observe that the structure-functional similarity between the empirical connectomes eFC and eSC used for the model derivation and validation exhibited weak opposite relationships between parameter clusters and across the number of the WBT streamlines as compared to the correspondence between simulated and empirical data (see S6 Fig a1 and b1 in Supplementary materials). This indicated a nontrivial character of the reported results that do not directly follow from the empirical structure-function correspondence.

It is interesting to note here that the best agreement between simulated and empirical functional data (sFC and eFC) was attained for the considered model at small (zero) delays (Fig 3). It is therefore safe to consider such a type of model simulating ultra slow BOLD dynamics without delay in coupling [35,36,44]. However, if other fitting modalities are involved, the delay in coupling can play an important role, for instance, for the structure-functional (sFC-eSC) model fitting. The latter can be used to mediate the impact of non-zero delay on the functional model validation, which also plays an important role for the higher correspondence between simulated and empirical functional data (Fig 6). The values of the optimal non-zero delays for the latter fitting modality can be influenced by the natural frequencies of oscillators (Eq 1) demonstrating relatively strong negative correlations with the structure-functional model fitting as illustrated in S7 Fig and S8 Fig in Supplementary materials. Consequently, the optimal speed of the signal propagation in the brain as revealed by the modeling results can be regulated by the mean intrinsic time scale of oscillatory activity of individual brain regions.

### Stratification analysis and the optimal condition

The problem of the optimal number of the total WBT streamlines was also addressed in this study beyond the group-level analysis and aimed at the best fitting of the personalized models for individual subjects. To investigate the impact of the WBT resolution at the level of individual subjects, we stratified the entire subject population into smaller subgroups with more homogeneous (heterogeneous) model dynamics within (between) subgroups. One of the stratification approaches is to show the effect of the graph-theoretical network properties modulated by the WBT density on performance of the model. We found that such correlations for individual subjects are well-pronounced for the Schaefer atlas, but they are relatively less expressed for the Harvard-Oxford atlas (Fig 5 a1 and b1). Nevertheless, the stratification can be designed by combining the splitting results for network properties, which resulted in a clear differentiation of the impact of the WBT streamline number on the model validation across stratified subgroups and brain parcellations (Fig 5).

Another approach to stratification of the subjects was based on the global delay parameter clustering for the structure-functional model fitting discussed above and can provide an informed view of the validation results for the functional model fitting (Fig 6). One more stratification approach is illustrated in Fig 7, where the subjects were split into two subgroups of qualitatively different individual behavior of the goodness-of-fit versus the streamline number. Based on the obtained results, we can propose to use the large number (~2M-10M) of the WBT streamlines for the best functional model validation, if the Schaefer atlas was used for the brain parcellation. On the other hand, the recommendation is completely opposite for more than 20% of subjects for the brain parcellation based on the Harvard-Oxford atlas (Fig 8 b2, blue dashed curve 3). For such subjects, the large number of streamlines can lead to a lower quality of the model fitting as compared to rather sparse WBT containing, for example, only 50K streamlines. Differentiating the subjects according to the discussed stratification criteria can help to design an individual data processing workflow and configurations of parameters for the optimal personalized modeling of the brain dynamics. In particular, based on the obtained results, we can suggest a personalized optimal number of the WBT streamlines for the considered brain parcellation for the better model performance at the modeling of the resting-state brain dynamics.

## Summary and conclusion

We found that varying number of total streamlines for WBT affects the network properties of the structural connectome and performance of the mathematical modeling of the resting-state brain dynamics. The results showed that a dense WBT is not always the best condition for the whole-brain mathematical modeling represented by a system of interacting oscillators with time delay in coupling. We also demonstrated that the optimal parameters of the data processing may be affected by the utilized brain parcellation that should be taken into account already at early steps of the data processing workflow. The present study did not aim to provide any quantitative conclusion concerning the optimal number of WBT streamlines, but rather to illustrate possible qualitative effects caused by the varying WBT density on the structural connectome and modeling results in combination with functional and anatomical brain parcellations. Our results can contribute to a better understanding of the interplay between the data processing and model parameters and their influence on data analytics of dwMRI and modeling of the resting-state fMRI data.

## Materials and methods

The current study considered 351 unrelated subjects (172 males, age 28.5 ± 3.5 years) from the Human Connectome Project (HCP) S1200 dataset [48]. HCP data (https://www.humanconnectome.org/, [48]) were acquired using protocols approved by the Washington University institutional review board (Mapping the Human Connectome: Structure, Function, and Heritability; IRB #201204036). Informed consent was obtained from subjects. Anonymized data are publicly available from ConnectomeDB (db.humanconnectome.org).

We reconstructed SC and PL by using six WBT densities and two atlases for individual subjects, then calculated simulated FC from simulated signals by the computational model composed of coupled phase oscillators with delayed coupling. We explored the two free parameters of the modeling for each subject and condition, and validated the model through the two model fitting modalities. We also calculated graph-theoretical network properties of SC and PL over considered conditions and compared the network properties with the maximal goodness-of-fit of the model. The individual subjects were stratified into groups of different network properties and modeling results.

### Preprocessing of MRI data and connectivity extraction

The current study used an *in-house* pipeline for the extraction of SC and PL matrices from the diffusion-weighted images (DWI). The pipeline consists of four modules: preprocessing of MRI and DWI data, WBT calculation, atlas transformation and connectivity reconstruction. It was optimized for parallel processing on high-performance computational clusters [49].

The pipeline was created with functions of Freesurfer [50], FSL [51], ANTs [52], and MRtrix3 [45]. Freesurfer was used for processing the T1-weighted image (T1) including bias-field correction, tissue segmentation, cortical (surface) reconstruction, volume-surface converting, and surface deformation for parcellation as well as for the correction of the eddy-current distortions and head-motion in DWIs using the corresponding b-vectors and b-values. MRtrix3 performed de-noising and bias-field correction on the DWIs. The pre-processed images were used for co-registration between the T1 and the DWIs and linear and non-linear transformation by functions of FSL. Linear and non-linear transformation matrices and images for registration from the standard MNI space to the native space and vice versa were estimated. Through the image registration, gray matter, white matter, cortical/subcortical, cerebellar, and cerebrospinal fluid masks were generated in the native DWI space.

The WBT calculation module included only MRtrix3 functions, where the response functions for spherical deconvolution were estimated using multi-shell-multi-tissue constrained deconvolution algorithm [53], fiber oriented distributions (FOD) were estimated from the DWIs using spherical deconvolution, and the WBT was created through the fiber tracking by the second-order integration over the FOD by a probabilistic algorithm [54]. In the latter step, we used six different numbers of total streamlines for varying WBT density: 10K, 50K, 100K, 500K, 2M, and 10M, where the “K” and “M” letters stand for thousand (Kilo-) and million (Mega-), respectively. The rest of the tracking parameters were set as default values of *tckgen* function from MRtrix3 documents [55].

The atlas transformation module applied the linear and non-linear transformation matrix and images to atlases that were sampled in the standard MNI space. We used Schaefer atlas with 100-area parcellation [46] and Harvard-Oxford atlas with 96 cortical regions [47]. After the transformation, the labeled voxels on the gray matter mask were selected for a seed and a target region. Consequently, the *tck2connectome* function of MRtrix3 reconstructed SC and PL (count and path-length matrices in Fig 1a).

For the empirical functional connectivity (FC), the BOLD signals were extracted from the resting-state fMRI data processed by ICA-FIX as provided by HCP repository [56]. The Schaefer atlas and the Harvard-Oxford atlas were applied for the parcellation of the processed fMRI into brain regions within the standard MNI 2 mm space (6th-generation in FSL). Empirical FC was calculated using Pearson’s correlation coefficient across BOLD signals extracted as mean signals of the parceled brain regions. There were four resting-state fMRI sessions (1200 volumes, TR = 720 ms) which consist of two different phase-encoding directions (left and right) scanned in different days. In addition, a concatenated BOLD signal was generated by using all four z-scored BOLD signals from the above four fMRI sessions, which results in five empirical FCs calculated for BOLD signals from the four fMRI sessions and the concatenated BOLD signals for each subject. Finally, 12 simulation conditions (6 WBTs ×2 atlases) were tested by simulation of the mathematical whole-brain model, where the model parameters were optimized for the best fit between simulated and empirical data.

### Mathematical whole-brain model

We simulated a whole-brain dynamical model of N coupled phase oscillators [19,32,38]

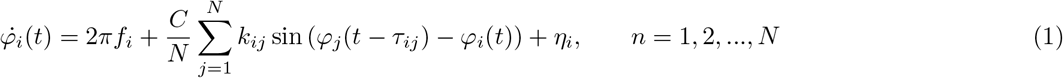

The number of oscillators *N* corresponds to the number of brain regions parceled as defined by a given brain atlas, where the phase *φ_i_(t)* models by sin (*φ_i_(t)*) the mean BOLD signal of the corresponding region. *C* is a global coupling which scales the level of couplings of the whole-brain network. *η_i_* is an independent noise perturbing of oscillator *i*, which is sampled from a random uniform distribution from the interval [−0.3,0.3]. The natural frequencies *f_i_* were estimated from the empirical data as frequencies of the maximal spectral peaks (restricted to the frequency range from 0.01 Hz to 0.1 Hz) of the empirical BOLD signals of the corresponding brain regions. *k_ij_* stands for the coupling strength between *i^th^* and *j^th^* oscillators, and *τ_ij_* approximates the time delay of the signal propagation between *i^th^* and *j^th^* oscillators. They were calculated from the empirical SC and PL and determined by the following equations:

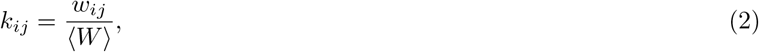

where *w_ij_* is the number of streamlines between *i^th^* and *j^th^* parceled regions and ⟨*W*⟩ is an averaged number of streamlines over all connections except self-connections, and

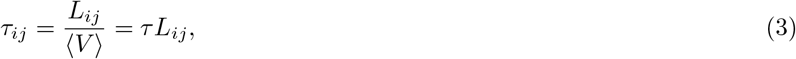

where τ is a global delay (unit: *s/m*) which is a reciprocal of an average speed of signal propagation ⟨V⟩ through the whole-brain network. The time step of the numerical integration of Eq 1 was fixed to 0.04 s, and the simulated signals were generated for 3500 seconds after skipping 500 seconds of the transient. The simulated BOLD signals and the corresponding simulated FCs were calculated from the phases downsampled to TR = 0.72 s, which is the repetition time of HCP fMRI.

The considered mathematical model Ep 1 has two main free parameters: the global coupling *C* and the global time delay τ. The global coupling ranged from 0 to 0.504 in evenly discretely distributed 64 values, and the global delay was from 0 to 423 s/m in evenly discretely distributed 48 values. Therefore, 3072 (64 ×48) simulations were performed for each subject to calculate the simulated FCs that were compared with empirical functional and structural data for each simulation condition. A total of 12,939,264 (64 ×48 ×12 ×351) simulations of model (Eq 1) were performed in this study for 351 subjects with 12 conditions (6 WBTs ×2 atlases). We explored the 2-dimensional (64 ×48) model parameter space and found the optimal parameter values for the best correspondence between simulated and empirical data. The correspondence was calculated by Pearson’s correlation coefficient between simulated FC (sFC) and empirical FC (eFC) and SC (eSC) depending on the model fitting modality. For each subject and simulation condition, 5 parameter planes of the *functional model fitting* modality (correlation between sFC and eFC as a goodness-of-fit of the model) were obtained corresponding to 5 eFCs. In addition, one parameter plane of the *structure-functional model fitting* modality (correlation between sFC and eSC) was also calculated. From each parameter plane, we selected the optimal (*C, τ*)-parameter point, where the maximal correlation between the simulated and the empirical data was reached.

### Effects of different WBT conditions

We revealed the effects of the varying WBT density on the modeling results by evaluating its impact on 1) the graph-theoretical network properties of empirical structural connectome, 2) patterns of the optimal model parameters in the model parameter space, and 3) model performance as given by the quality of the model fitting over simulation conditions. Based on the results from the three approaches, we introduced three criteria (see below) for differentiation of the influence of the WBT density on the modeling results for individual subjects. To do this, we stratified the entire subject population by splitting it into several subgroups according to the mentioned criteria based on *(i)* the relationships between the network properties and the results of the functional model fitting over WBT conditions, *(ii)* distributions of the optimal model parameters of the structure-functional model fitting, and *(iii)* positive and negative slopes (increments) of the maximal goodness-of-fit (the model performance) values across the two extreme cases of the considered 10K and 10M WBT streamlines for individual subjects.

### Structural architecture and network properties over WBT conditions

To investigate the impact of the varying WBT density on the architecture of structural networks, we calculated graph-theoretical network properties from SC and PL for each subject, WBT condition, and atlas. The considered 10 network properties (8 local properties and 2 global properties) included the weighted node degree, clustering coefficient, betweenness centrality, local efficiency, global efficiency, and modularity, which were calculated by the brain connectivity toolbox in Matlab [57]. For the local properties, both the average (Avg.) and the standard deviation (S.D.) were calculated. For the considered 6 WBT conditions and eFC of the concatenated session, we obtained the optimal goodness-of-fit of the model (the functional model fitting), which were correlated with a given network property across 6 WBT conditions for each individual subject. Consequently, based on the obtained 10 distributions of the correlation coefficients for all subjects, we split the subjects into two subgroups: 1) with positive correlations between the network properties and the maximal functional goodness-of-fit and 2) with negative correlations. We then compared the results of the functional model fitting between the two subject subgroups for every of the considered network properties and WBT conditions by the two-sample one-tail t-test to evaluate which subgroup shows significantly better functional model fitting than the other, see S5 Fig in Supplementary materials. Based on the results of such a comparison, we selected the network properties and the corresponding subject subgroups with significantly higher model performance for 10M, 2M, 500K, or 100K WBT streamlines. Then the subjects in an intersection of the selected subgroups constituted a group referred to as pattern 1, and the rest of them were referred to as pattern 2.

### Impact of time delay on the model fitting

For the second stratification criterion, the optimal model parameters of the maximal correspondence between sFC and eSC were divided into two clusters as suggested by the their bimodal distribution splitting small and large values of the optimal time delay (Fig 6). Since subjects can move between the parameter clusters when the total number of the WBT streamlines varies from 10M to 10K, we separated the subjects into five classes: Always staying in cluster 1 (From 1 to 1) or in cluster 2 (From 2 to 2), only once moving either from cluster 1 to cluster 2 (From 1 to 2) or in opposite direction (From 2 to 1), and performing multiple changing between the two clusters (Multiple). This approach based on the distribution of the optimal model parameters was used as the second criterion for the stratification of subjects.

### Variation of the model performance

The last stratification criterion is based on the behavior of the optimal goodness-of-fit values when the number of WBT streamlines varies. To quantify it, we calculated the increment of the maximal similarity between sFC and eFCs matrices of the concatenated session for every individual subject when the number of the WBT streamlines increases from 10K to 10M. Then, all subjects were divided into two subgroups exhibiting either positive or negative slopes (increments) of the goodness-of-fit behavior versus the number of WBT streamlines (Fig 7). According to this criterion, the subject were stratified into two subgroups demonstrating the best functional model fitting for either maximal or minimal number of the WBT streamlines considered. Consequently, we used all three criteria for the three-step stratification analysis (Fig 8).

## Acknowledgments

The authors are grateful to Esther Florin for fruitful discussions, helpful comments on the manuscript and advices concerning statistical analysis. The authors also gratefully acknowledge the computing time granted through JARA on the supercomputer JURECA at Forschungszentrum Jülich. This work was supported by the Portfolio Theme Supercomputing and Modeling for the Human Brain by the Helmholtz association (https://www.helmholtz.de/en), the Human Brain Project, and the European Union’s Horizon 2020 Research and Innovation Programme (https://cordis.europa.eu) under Grant Agreements 720270 (HBP SGA1), 785907 (HBP SGA2), 945539 (HBP SGA3), and 826421 (VirtualBrainCloud). The funders had no role in study design, data collection and analysis, decision to publish, or preparation of the manuscript.

Data were provided S1200 by the Human Connectome Project, WU-Minn Consortium (Principal Investigators: David Van Essen and Kamil Ugurbil; 1U54MH091657) funded by the 16 NIH Institutes and Centers that support the NIH Blueprint for Neuroscience Research; and by the McDonnell Center for Systems Neuroscience at Washington University.

## Supplementary materials

**S1 Fig.**
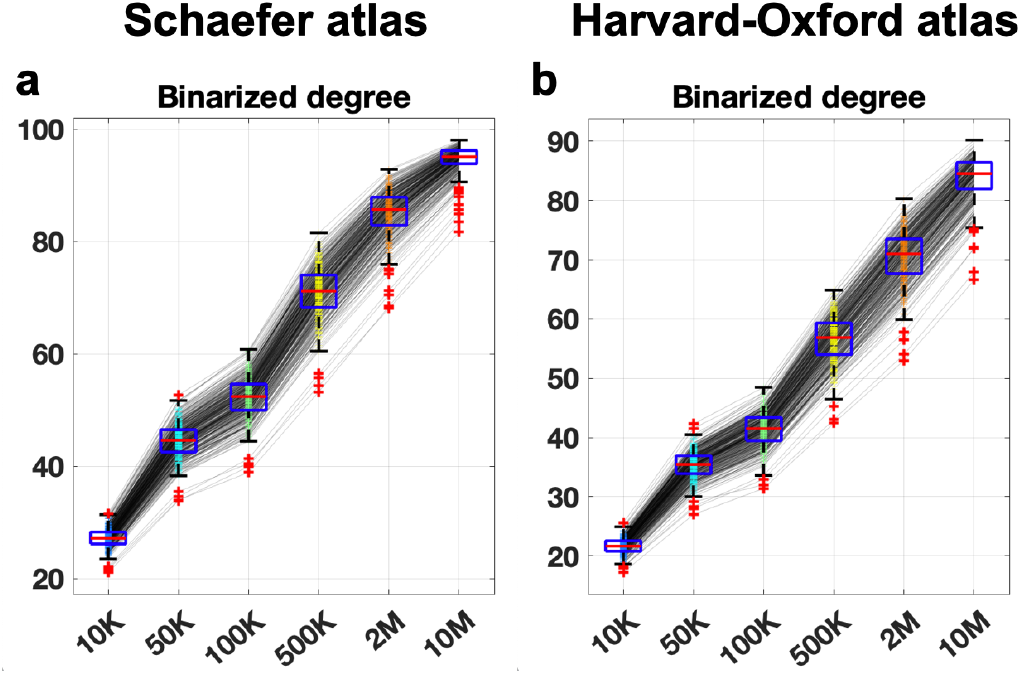
Average binarized node degrees over the WBT conditions for **(a)** the Schaefer atlas and **(b)** the Harvard-Oxford atlas.

**S2 Fig.**
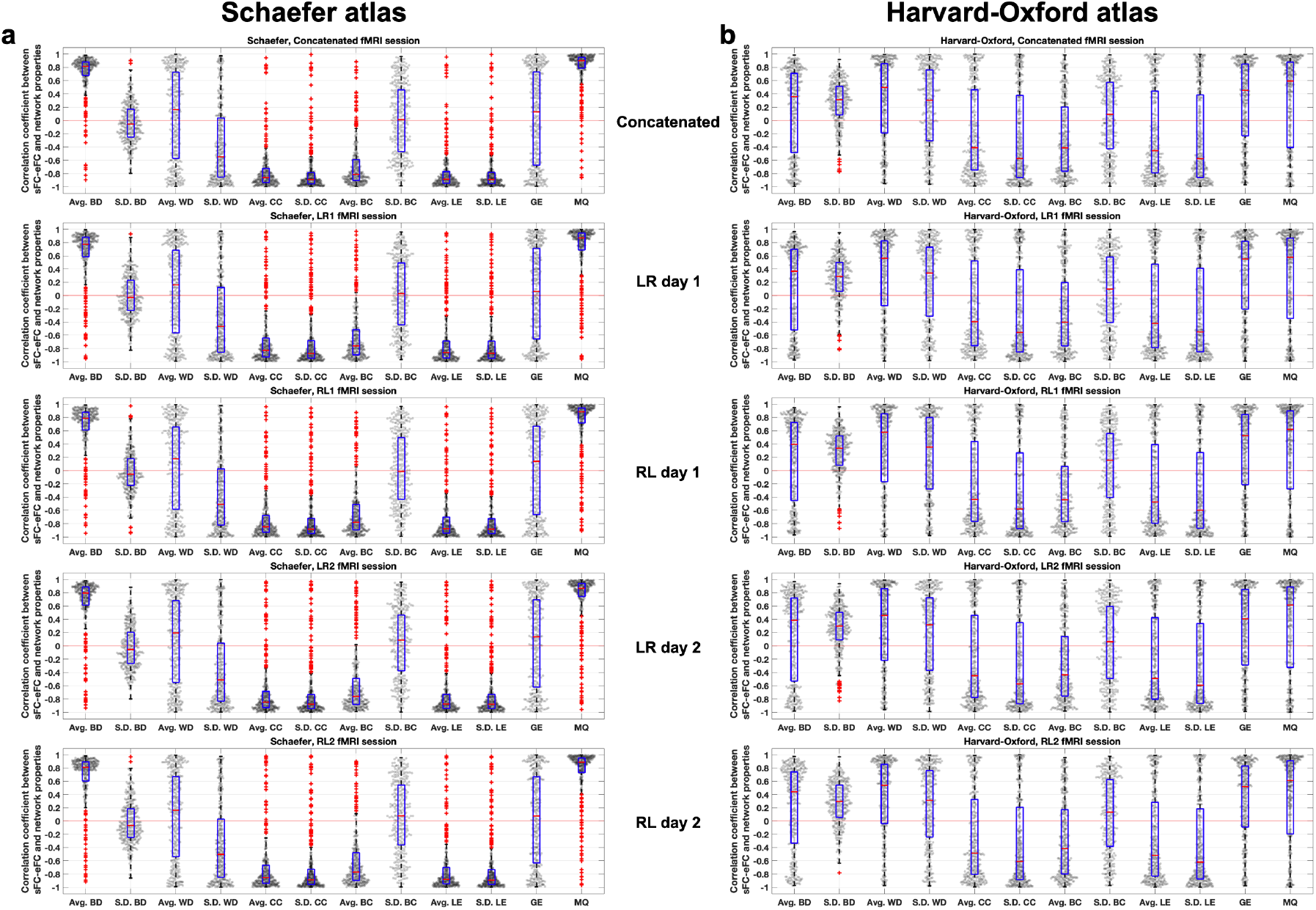
Relationships between the network properties and the results of the functional model fitting (maximal similarity between sFC and eFC) for individual subjects and 5 sessions for **(a)** the Schaefer atlas and **(b)** the Harvard-Oxford atlas. The gray dots represent the values for individual subjects, and the box plots illustrate the medians (red lines), the interquartile ranges (blue boxes) and the outliers (red pluses). Abbreviations of the network property names: binarized node degree (BD), weighted node degree (WD), clustering coefficient (CC), betweenness centrality (BC), local efficiency (LE), global efficiency (GE), and modularity Q (MQ).

**S3 Fig.**
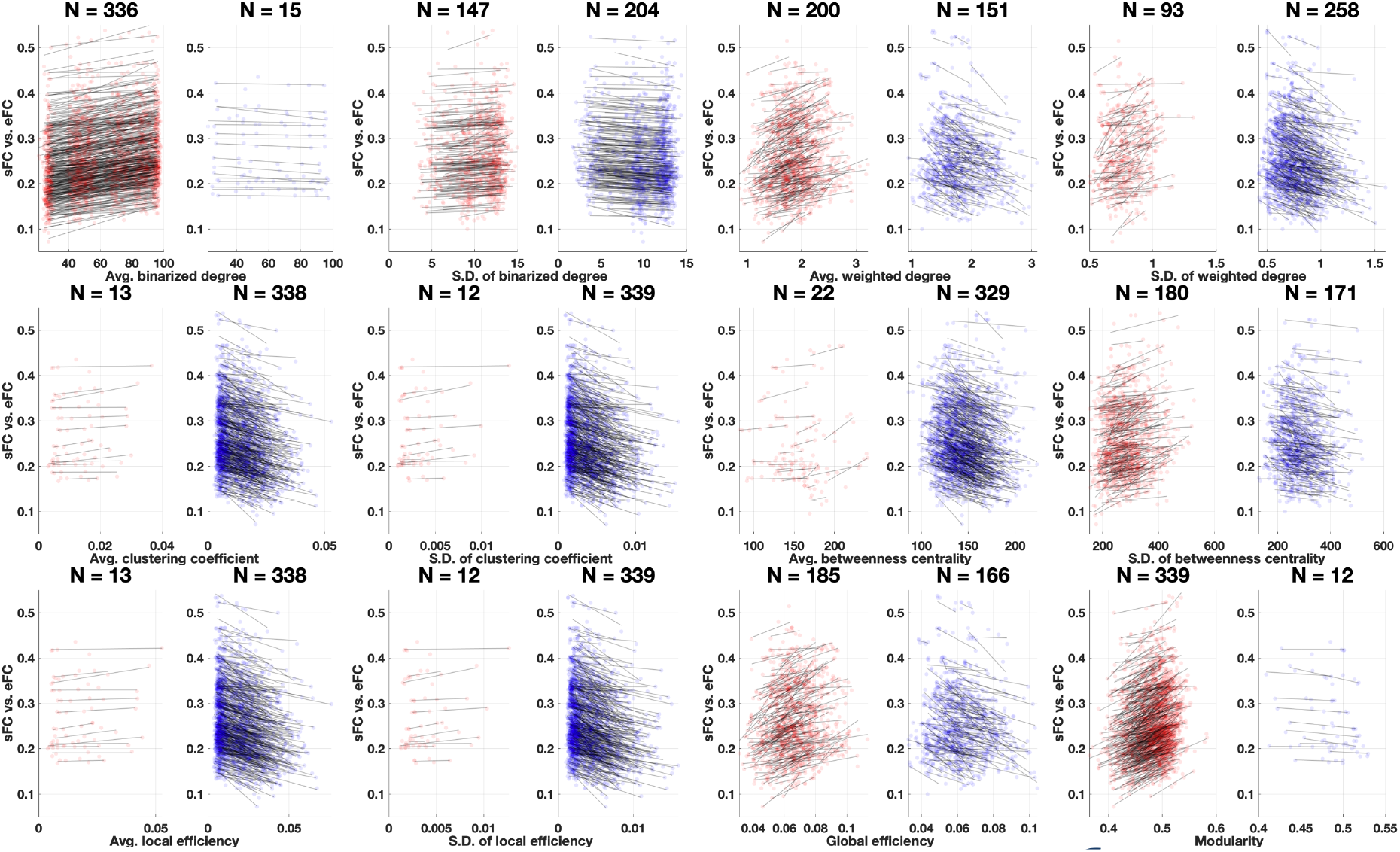
Pearson’s correlation coefficients between network properties and the goodness-of-fit of the model (similarity between sFC and eFC) for the Schaefer atlas.

**S4 Fig.**
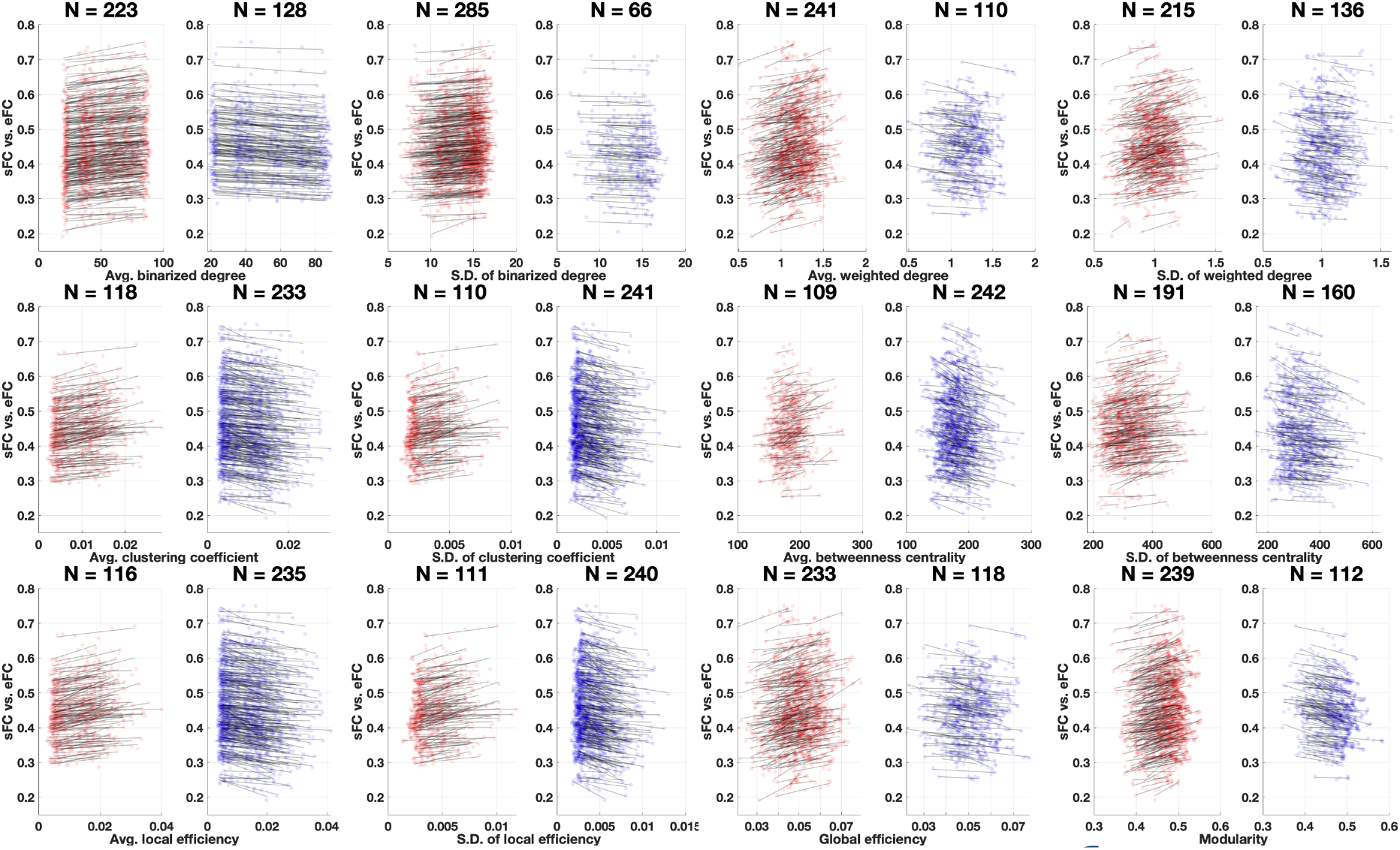
Pearson’s correlation coefficients between network properties and the goodness-of-fit of the model (similarity between sFC and eFC) for the Harvard-Oxford atlas.

**S5 Fig.**
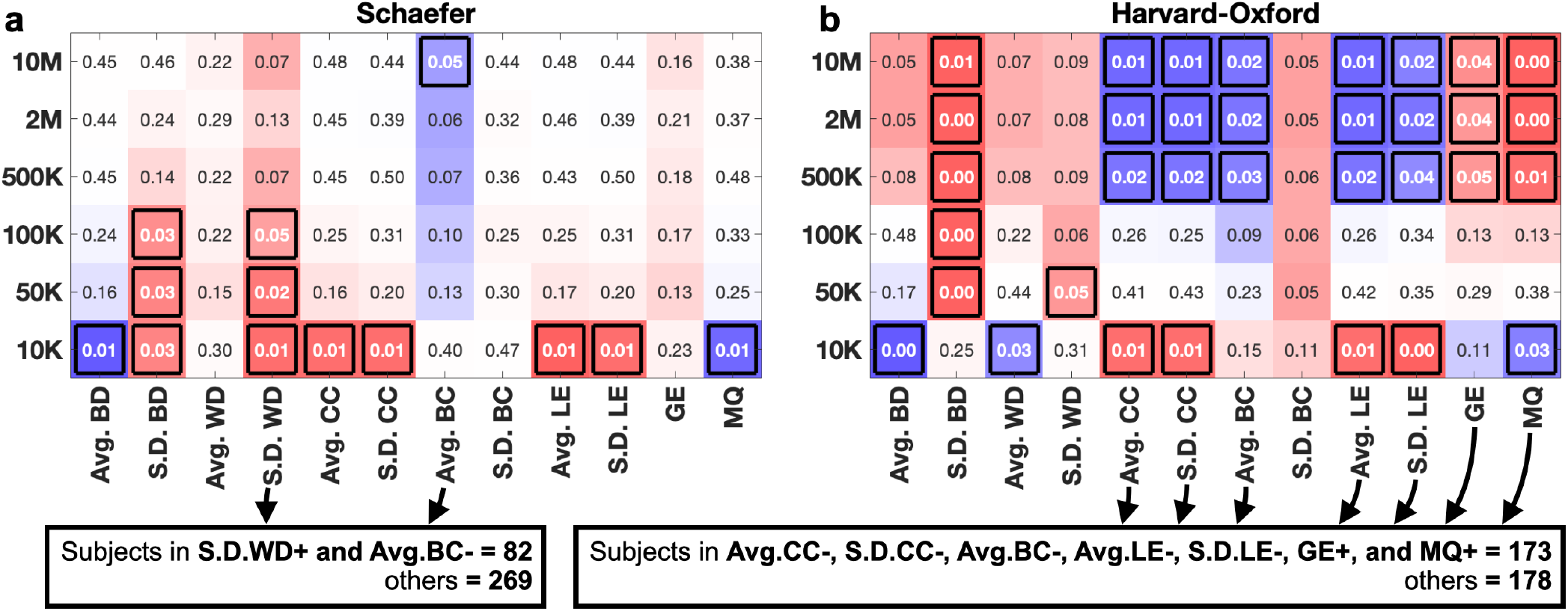
Stratification of subjects based on the distributions of the correlation coefficients between the network properties and the maximal functional goodness-of-fit values of the model for **(a)** the Schaefer atlas and **(b)** the Harvard-Oxford atlas. **(a, b)** p-values of the two-sample one-tail t-test for the differences of the model fitting between the two subject subgroups split based on the positive or negative values of the correlation coefficients whose distributions are illustrated in Fig 5 a1 and b1. The corresponding network properties and the number of WBT streamlines are indicated on the horizontal and vertical axes, respectively. The black squares in the tables indicate significant results with *p* < 0.05. Red cells mean the subgroup of positive correlations showed higher goodness-of-fit of the model than the subgroup of negative correlations. Blue cells mean the subgroup of negative correlations showed higher goodness-of-fit of the model than the subgroup of positive correlations. Abbreviations of the network property names are as in Fig 5. The plus (+) or minus (−) sign after the property name in the black boxes, e.g., **S.D. WD+**, indicates a group of subjects with positive, respectively, negative correlation coefficients between the corresponding network property and the model goodness-of-fit values.

**S6 Fig.**
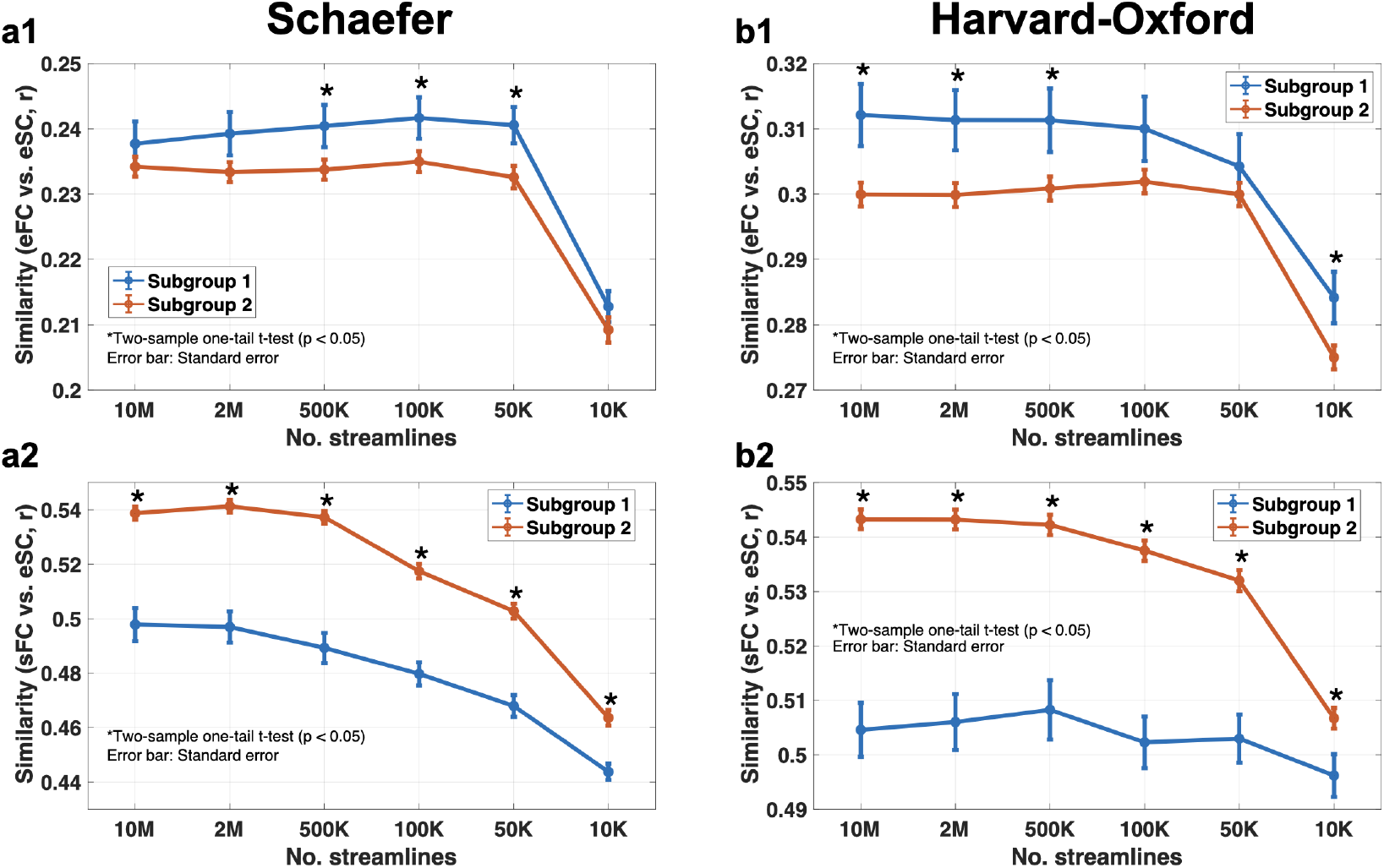
**(a1 and b1)** Agreements between eFC and eSC for two subgroups by the second criterion of the stratification. (a2 and b2) Agreements between sFC and eSC for two subgroups by the second criterion of the stratification. The subgroup 1 means the cluster 1 and the subgroup 2 means the cluster 2 in Fig 6.

**S7 Fig.**
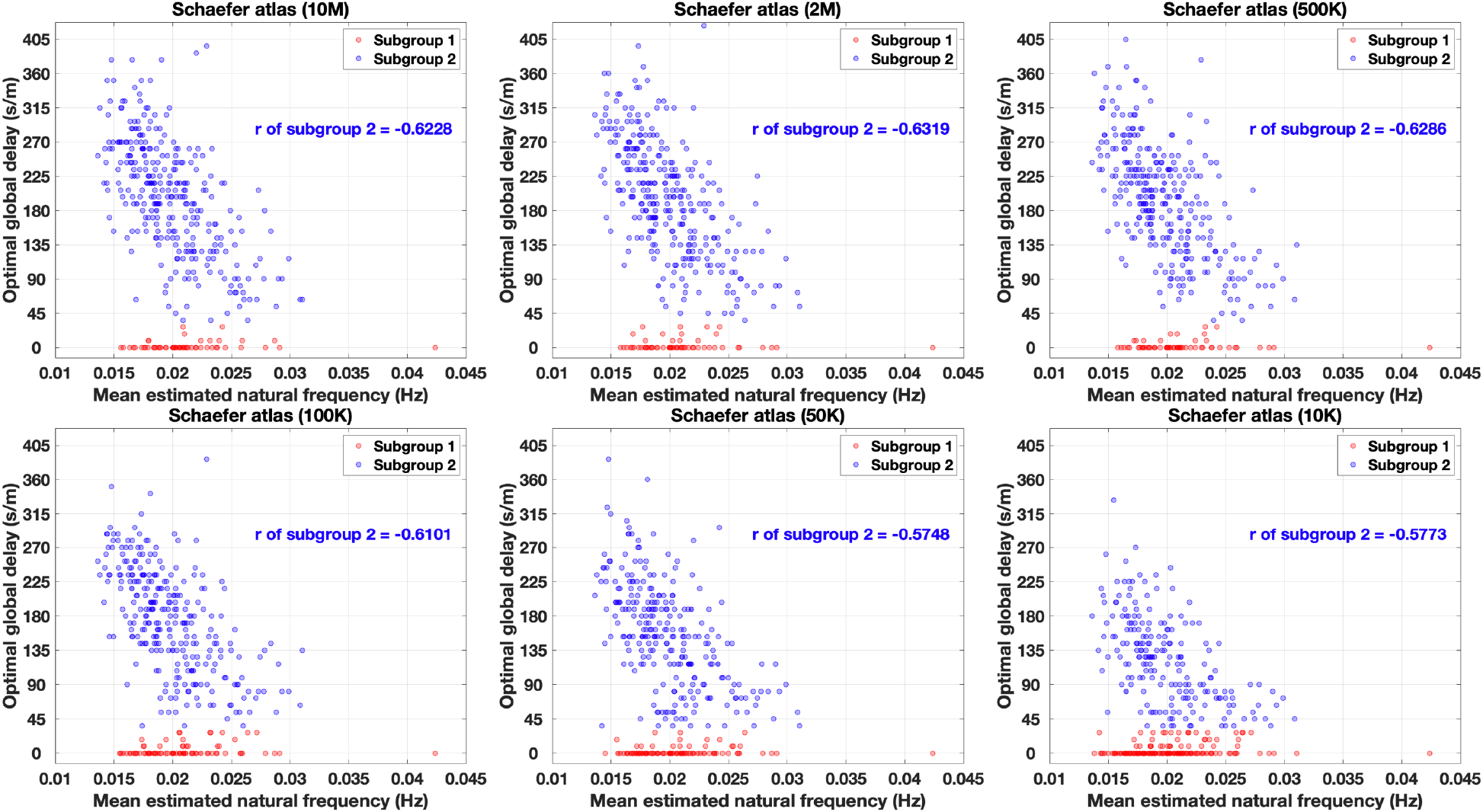
Relations between the mean estimated natural frequencies and the optimal global delays for the similarities of sFC versus eSC for the Schaefer atlas. Red dots are the optimal parameter sets of subgroup 1 and blue dots are that of subgroup 2 in Fig 6. The subgroup 1 means the cluster 1 and the subgroup 2 means the cluster 2 in Fig 6.

**S8 Fig.**
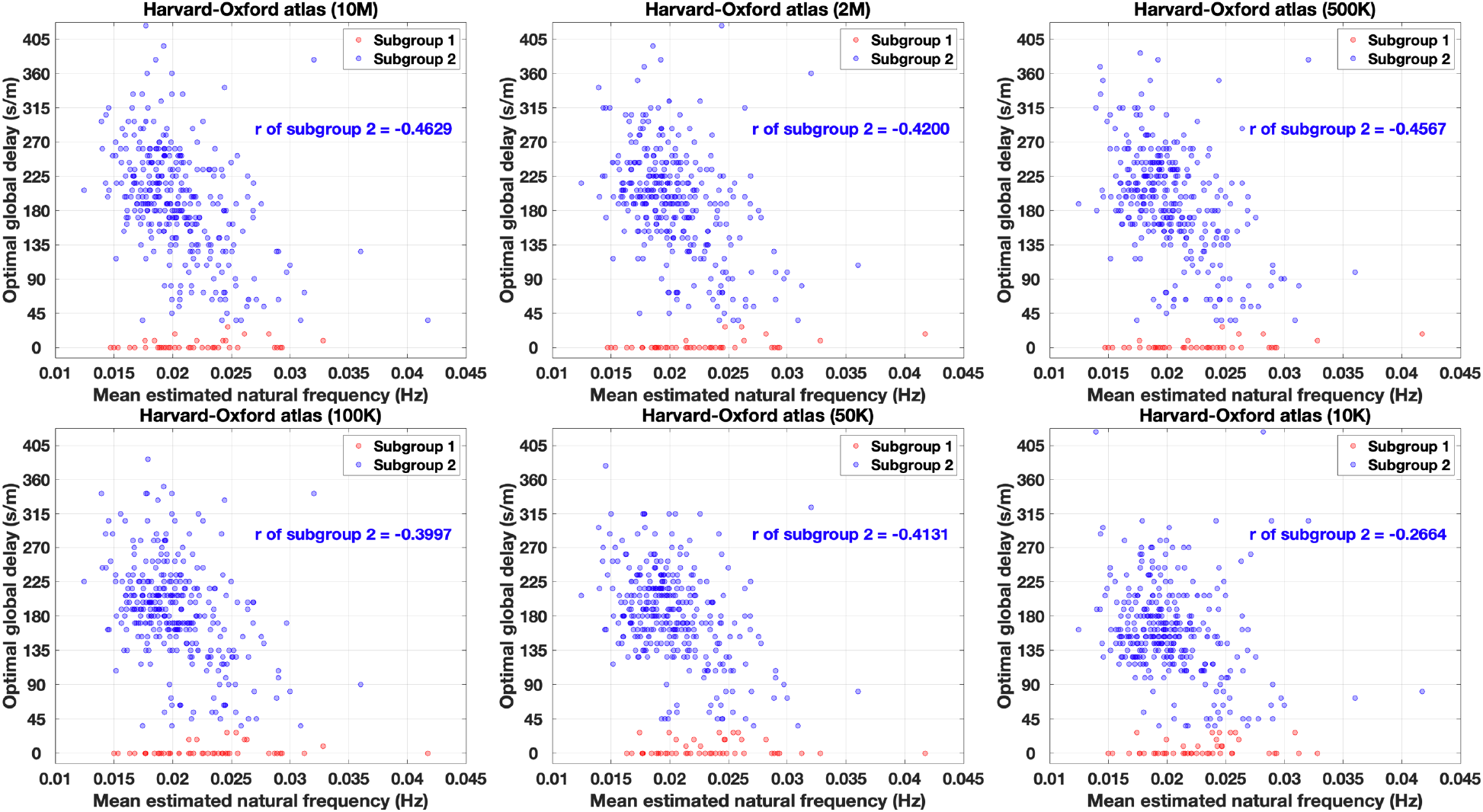
Relations between the mean estimated natural frequencies and the optimal global delays for the similarities of sFC versus eSC for the Harvard-Oxford atlas. Red dots are the optimal parameter sets of subgroup 1 and blue dots are that of subgroup 2 in Fig 6. The subgroup 1 means the cluster 1 and the subgroup 2 means the cluster 2 in Fig 6.

